# Detecting the effect of genetic diversity on brain composition in an Alzheimer’s disease mouse model

**DOI:** 10.1101/2023.02.27.530226

**Authors:** Brianna Gurdon, Sharon C. Yates, Gergely Csucs, Nicolaas E. Groeneboom, Niran Hadad, Maria Telpoukhovskaia, Andrew Ouellette, Tionna Ouellette, Kristen O’Connell, Surjeet Singh, Tom Murdy, Erin Merchant, Ingvild Bjerke, Heidi Kleven, Ulrike Schlegel, Trygve B. Leergaard, Maja A. Puchades, Jan G. Bjaalie, Catherine C. Kaczorowski

**Affiliations:** The Jackson Laboratory, Bar Harbor, ME; The University of Maine Graduate School of Biomedical Sciences and Engineering, Orono, ME; Neural Systems Laboratory, Institute of Basic Medical Sciences, University of Oslo, Oslo, Norway; Tufts University Graduate School of Biomedical Sciences, Medford, MA

**Keywords:** mouse brain, reference atlas, immunohistochemistry, deconvolution, Alzheimer’s disease, neurodegeneration, cell composition, genetic diversity

## Abstract

Alzheimer’s disease (AD) is characterized by neurodegeneration, pathology accumulation, and progressive cognitive decline. There is significant variation in age at onset and severity of symptoms highlighting the importance of genetic diversity in the study of AD. To address this, we analyzed cell and pathology composition of 6- and 14-month-old AD-BXD mouse brains using the semi-automated workflow (QUINT); which we expanded to allow for nonlinear refinement of brain atlas-registration, and quality control assessment of atlas-registration and brain section integrity. Near global age-related increases in microglia, astrocyte, and amyloid-beta accumulation were measured, while regional variation in neuron load existed among strains. Furthermore, hippocampal immunohistochemistry analyses were combined with bulk RNA-sequencing results to demonstrate the relationship between cell composition and gene expression. Overall, the additional functionality of the QUINT workflow delivers a highly effective method for registering and quantifying cell and pathology changes in diverse disease models.

## Introduction

Alzheimer’s Disease (AD) is a multifaceted neurodegenerative condition that currently has no cure and impacts millions around the globe^1^. AD is characterized by the accumulation of amyloid-beta (AB) plaques, neurofibrillary tau tangles, severe gliosis, and progressive neurodegeneration^2^, leading to clinical symptoms and cognitive decline that eventually lead to death^3^. There is significant variation in the age at symptom onset and severity of cognitive decline, with highly susceptible individuals exhibiting early onset and rapid decline, while resilient individuals remain cognitively intact late in life^4, 5^. Further characterization of pathology development including neurodegeneration, amyloid-beta deposition, and neuroinflammation is needed to better understand the impact of this variation on clinical disease outcomes. Moreover, this characterization is highly relevant since changes in the composition of brain tissue and the development of neuropathology can precede (and might even predict) clinical symptoms, and therefore serves as a valuable resource for defining disease subtypes and possible mechanisms of resilience^6–8^.

Mouse models of AD offer the opportunity to study changes in brain pathology in a controlled manner to gain a better understanding of how AD manifests and may progress in humans^9, 10^. In these models, organism-wide, brain-wide, or region-specific imaging and omics approaches can be implemented for the investigation of disease stages using cross-sectional or longitudinal study designs. To combat the lack of heterogeneity of traditional inbred AD mouse models, the AD-BXD mouse population that better recapitulates the complex heterogeneity of genetic, molecular, and cognitive features of human aging and AD was utilized in this study^11, 12^. The AD-BXD population was generated by crossing the C57BL/6J(B6)-5XFAD AD mouse model with strains from the BXD panel^11^. Despite being driven by alleles typically found in cases of early-onset AD, in the genetically diverse BXD strains, the 5XFAD transgene leads to a spectrum of phenotypes that recapitulate the clinical and pathological variation of late-onset AD^11, 13–16^. Since the relationship between symptomatology and changes in the composition of brain tissue is not fully understood, assessing changes in cell and pathology organization across a mouse population that models the heterogeneity of human AD may highlight brain regions and cell types associated with cognitive susceptibility or resilience to neurodegeneration, gliosis, and pathology^17–23^.

In addition to characterizing AD with imaging outcomes of cell composition in mouse models, changes with AD can be described by investigating deviations in gene expression among different cell types of the brain. Bulk RNA-sequencing (RNAseq) is a common method to study gene expression profiles of brain regions of interest; however, it is crucial to note that gene expression data generated from a tissue sample reflects an average gene expression profile across heterogeneous populations of cells^24^. Consequently, consideration of individual differences in regional cell composition is vital when interpreting the results from RNAseq data from different mouse strains and patient samples. Since AD has a substantial impact on brain structure, observed changes in gene expression in bulk tissue are likely to be masked by changes in cell-type composition across varying disease stages. In many AD studies that conduct RNAseq to determine disease signatures, it is not clear whether observed differences in gene expression among AD samples or between AD samples and controls are due to changes in transcriptional regulation or the relative proportions of different cell types in the tissue samples^14, 25, 26^. Measuring cell composition and recognizing the contribution of cell abundance when associating gene expression to disease traits is important for reducing spurious associations between AD phenotypes and gene expression^27, 28^. Deconvolution methods have been created in an attempt to estimate the proportions of different cell types in RNAseq results and to distinguish changes in gene expression stemming from changes in cell-type compositions versus alterations in gene activity^29–34^; however, the performance of deconvolution tools are highly variable^27, 35^.

Immunohistochemistry (IHC) quantification is the gold standard for measuring the cell composition of a tissue sample. When combined with brain-wide analysis methods that utilize reference atlases of the brain^36, 37^, IHC is a powerful tool that can be used to better understand the changes in cell composition that occur with age and AD, and the relative relationship between cellular load and gene expression. The QUINT workflow^38^ is one such semi-automated analysis method that combines a tool for registering histological brain section images (QuickNII^39^) to a reference atlas of the brain, with tools for extracting (ilastik^40^) and quantifying IHC-stained features (Nutil^41^). A key step in the QUINT workflow is that customized atlas-plates, derived from a three-dimensional brain atlas, are linearly registered to brain section images^39^. However, with morphological differences seen among mouse strains, disease states, and ages^42–46^, and morphological distortions occurring during histological processing, linear registration is often insufficient to achieve accurate anatomical registration. This motivated the expansion of the QUINT workflow with new functionality to increase the quality of the atlas-registration by application of nonlinear refinements (VisuAlign); as well as providing a means to verify the atlas-registration by systematic random sampling (QCAlign). Here, we utilize the expanded QUINT workflow to characterize regional composition of neurons, reactive astrocytes, microglia, and amyloid beta pathology across brains of AD-BXD mice at different ages, in regions defined by the Allen Mouse Brain Common Coordinate Framework v3 (CCFv3). By completing this analysis, we provide an expansive brain-wide characterization of diverse 5XFAD mice and 1). assess changes in cell and pathology composition between AD-BXD animals at 6 and 14-months of age, 2). assess variation in cellular abundance among AD-BXD strains, and 3). interpret bulk RNAseq data with respect to the cellular-abundance, in order to differentiate effects driven by AD from effects driven by cellular composition in the hippocampal formation.

## Methods

### Method relating to the mice and IHC

#### Bioethics

All mouse experiments occurred at the University of Tennessee Health Science Center and were carried out in accordance with the principles of the Basel Declaration and standards of the Association for the Assessment and Accreditation of Laboratory Animal Care (AAALAC), as well as the recommendations of the National Institutes of Health Guide for the Care and Use of Laboratory Animals. The protocol was approved by the Institutional Animal Care and Use Committee (IACUC) at the University of Tennessee Health Science Center.

#### Animals

All data used in this study are from the AD-BXD panel, which have been previously described^11^ (Figure 1b). Briefly, female B6 mice hemizygous for the 5XFAD transgene (B6.Cg-Tg(APPSweF1LonPSEN1*M146L*L286V)6799Vas/ Mmjax, Stock No. #24848-JAX) were mated to males from the BXD genetic reference panel resulting in sets of isogenic F1 AD-BXD strains that either harbor the 5XFAD transgene or are nontransgenic (Ntg)-BXD littermate “normal aging” controls. Male and female AD-BXD mice were group housed as a mix of 5XFAD and Ntg same-sex littermates (2-5 per cage) and maintained on a 12-hour light–dark cycle with *ad libitum* access to food and water. All mice were genotyped for the 5XFAD transgene through a combination of in-house genotyping according to The Jackson Laboratory Transgenic Genotyping Services protocols for strain #34848-JAX and outside services (Transnetyx, TN, USA). This study included a total of 40 mice (2 males and 38 females) of 6 months (6m; n=20) and 14 months (14m; n=20). These included 29 mice from 14 AD-BXD strains (n = 1-4 mice per strain); 8 mice from founder strains C57BI/6J (B6) 5XFAD (n = 2), and F1 B6/DBA/2J (D2) 5XFAD (n = 6); and 3 Ntg-BXD mice (all 6 m). An overview of all the animals included in the study is given in Supplementary Table 1.

**Figure 1.**
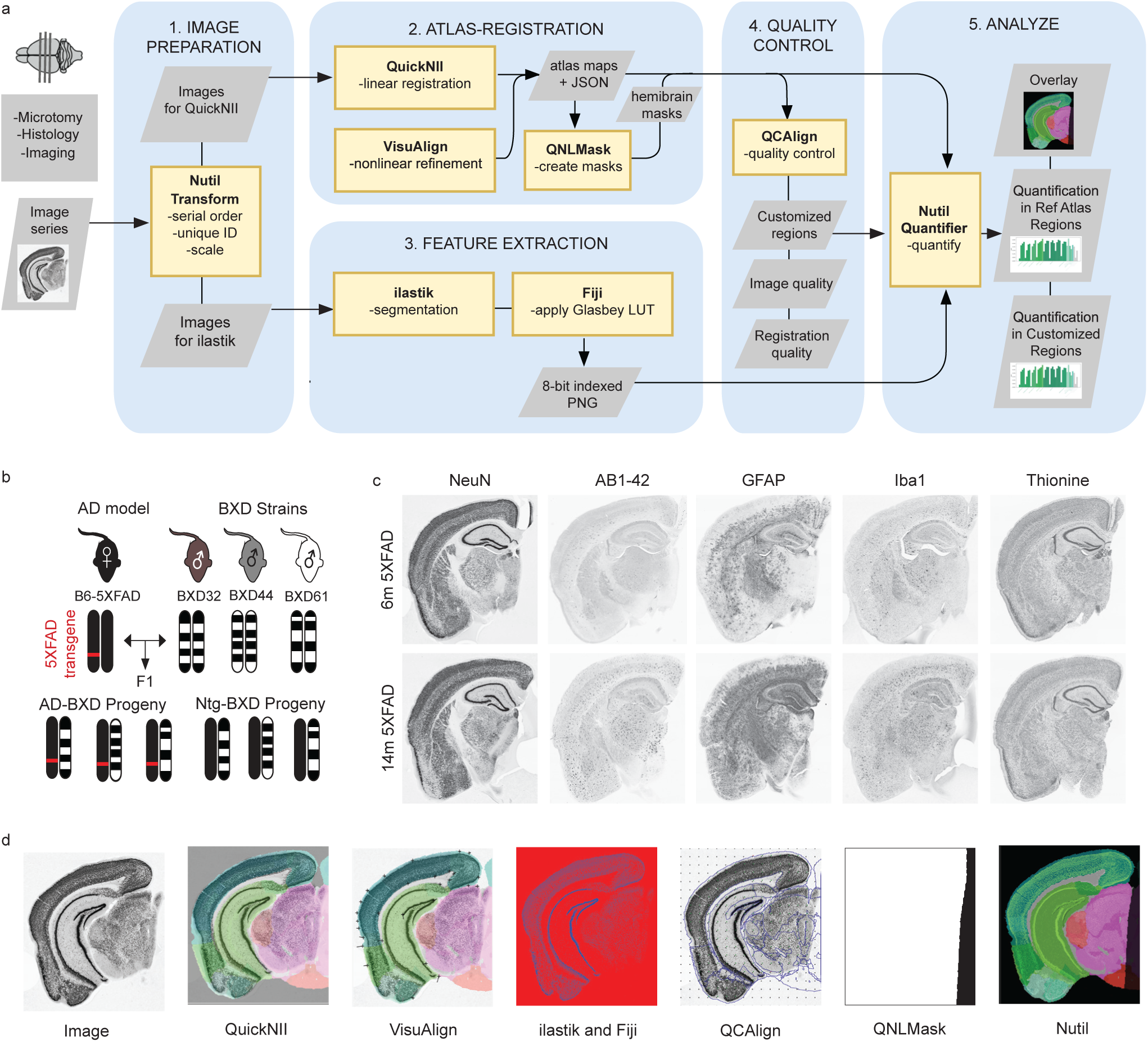
Study design and QUINT workflow overview. **a.)** Regional pathology and cell composition were quantified using the expanded QUINT workflow. 1) Raw images were processed to meet size requirements. 2.) Brain sections were registered to the Allen Mouse Brain Atlas CCFv3 2015 in QuickNII and refined using VisuAlign. Hemibrain masks were created in QNLMask 3.) Ilastik pixel classification was used to establish cell detection parameters for each stain and converted to RBG format in FIJI. 4.) Post-registration quality control assessment was performed using the novel QCAlign tool. 5.) Segmentation, registration, and mask creation steps were combined using Nutil to receive percent stain-positive cell coverage per region area. **b.)** Immunohistochemistry was completed for an experimental cohort of 40 mice from the AD-BXD mouse model of AD (see Supplemental Table 1). Adapted from Neuner et al., 2019. **c.)** Brain sections of 6m and 14m mice were sectioned and stained for thionine, NeuN, GFAP, Iba1, and AB1-42 via Neuroscience Associates. **d.)** Representative images from each step in the QUINT workflow.

#### Immunohistochemistry

##### Tissue collection and shipment

Mice were deeply anesthetized using isoflurane before decapitation and rapid removal of the brain at appropriate time points (6m or 14m). The hypothalamus was dissected out and the brain was bisected down the sagittal midline. One half of the brain was immediately further dissected and snap frozen to be used for RNAseq and the other hemisphere was placed in 4% paraformaldehyde and kept at 4°C to be used for IHC as previously described^11, 13, 16^. In order to minimize technical variation in IHC, hemibrains were sent overnight to Neuroscience Associates (Knoxville, TN), where the cerebellum was removed and hemibrains were embedded, processed, and stained simultaneously in blocks of 40.

##### Neurohistology Embedding and Sectioning

Hemibrains received at Neuroscience Associates were examined for overall tissue integrity (no major damage or tissue breakdown), then treated overnight with 20% glycerol and 2% dimethylsulfoxide to prevent freeze-artifacts. The specimens were then embedded in a gelatin matrix using MultiBrain®/ MultiCord® Technology (Neuroscience Associates, Knoxville, TN). The blocks were rapidly frozen, after curing by immersion in 2-Methylbutane chilled with crushed dry ice and mounted on a freezing stage of an AO 860 sliding microtome. The MultiBrain®/ MultiCord® blocks were sectioned in coronally with desired micrometer (40µ) setting on the microtome. All sections were cut through the entire length of the specimen and collected sequentially into series of 24 containers. All containers contained Antigen Preserve solution (50% PBS pH7.0, 50% Ethylene Glycol, 1% Polyvinyl Pyrrolidone); no sections were discarded.

##### IHC staining

Free floating sections were stained for Aβ1-42 (amyloid beta pathology), glial fibrillary acidic protein (GFAP, reactive astrocytes) and ionized calcium binding adapter protein 1 (Iba1, microglia) on every 24th section spaced at 960 μm, yielding approximately 9 sections per hemibrain. Staining for NeuN (neurons) and thionine (Nissl, cell bodies) was performed on every 12th section spaced at 480 μm, yielding approximately 19 sections per hemibrain. For Aβ1-42, GFAP, Iba1 and NeuN, all incubation solutions from the blocking serum onward used Tris buffered saline (TBS) with Triton X-100 as the vehicle; all rinses were with TBS. After a hydrogen peroxide treatment and blocking serum, the sections were immunostained with the primary antibodies, as shown in Supplemental Table 2, overnight at room temperature. Vehicle solutions contained Triton X-100 for permeabilization. Following rinses, a biotinylated secondary antibody was applied. After further rinses Vector Lab’s ABC solution (avidin-biotin-HRP complex; VECTASTAIN® Elite ABC, Vector, Burlingame, CA) was applied. The sections were again rinsed, then treated with diaminobenzidine tetrahydrochloride (DAB) and hydrogen peroxide to create a visible reaction product. Following further rinses, the sections were mounted on gelatin-coated glass slides and air dried. The slides were dehydrated in alcohols, cleared in xylene and cover slipped. For thionine-Nissl Staining sections were mounted on gelatin-coated glass slides, air dried and carried through the following sequence: 95% ethanol, 95% ethanol/Formaldehyde; 95% ethanol, Chloroform/Ether/absolute ethanol (8:1:1), 95% ethanol; 10% HCl/ethanol, 95% ethanol, 70% ethanol, deionized water, thionine (0.05% thionine/acetate buffer, pH 4.5) (Fisher, T40925), deionized water, 70% ethanol, 95% ethanol, Acetic Acid/ethanol, 95% ethanol, 100% ethanol, 100% ethanol, 1:1 100% ethanol/xylene, xylene, xylene, coverslip.

##### Slide identification and imaging

Each slide was laser etched with the block number and the stain. Following serial ordering of the slides, rostral to caudal for each stain, the slides were numbered by permanent ink in the upper right corner.

Neuroscience Associates (NSA) performed scanning of each slide at 20x using a Huron Digital Pathology TissueScope LE120 (0.4 microns/pixel). Brain image series were compiled by reconstructing the IHC sections as sliced and indicated by NSA.

Further information and requests for resources and reagents should be directed to and will be fulfilled by the Corresponding author.

### Methods Related to QUINT Workflow Utilization

#### QUINT workflow development

The QUINT workflow supports brain-wide quantification of IHC data in relation to a reference atlas such as the Allen Mouse Brain Common Coordinate Framework v3 (CCFv3). In the workflow (Figure 1a), the QuickNII software^39^ is used to spatially register atlas-plates from a 3D digital brain atlas to serial section images, the ilastik software^40^ is used to extract features from the images, and the Nutil software^41^ is used to quantify features per atlas-region. To meet the needs of the current project, two new software, VisuAlign (RRID: SCR_017978) and QCAlign (RRID:SCR_023088), were developed and integrated in the QUINT workflow. VisuAlign is used to apply in-plane nonlinear refinements of the atlas to achieve the best fit over the section images. This task is performed by visually identifying mismatches between section images and the corresponding atlas plates, and manually assigning a set of anchor points denoting corrections. VisuAlign then uses these anchor points to create a continuous, nonlinear deformation field covering the entire section image. QCAlign is used to 1. detect sections or regions not suited for QUINT analysis (i.e., due to damage), and 2. to assess the quality of the atlas-registration to each region in the sections. Both QCAlign assessments are performed by systematic random sampling. The second assessment is based on anatomical expertise by evaluating how well delineations supplied by the atlas match up with boundaries revealed by IHC-staining. Since validation of the atlas-registration is only possible for regions that have visible boundaries in the sections, and reference atlases are structured in systematic hierarchies that group related regions^47^, functionality was also implemented in QCAlign for adjusting the hierarchy to a customized level that supports verification of the regional registration (i.e. a level where the delineations from the atlas roughly matching the boundaries that are visible in the sections). This customized hierarchy level can be exported as a TXT file and used in the Nutil software to define customized regions to use for the brain-wide quantification.

#### Image Pre-processing

To perform stain segmentation in ilastik, the images were inspected, cropped, and downscaled using different scaling factors for the different stains (AB1-42: 0.20, GFAP: 0.40, Iba1: 0.40, NeuN: 0.40, thionine: 0.35). Scaling factors were determined by gradually increasing the scaling factor and manually determining the level at which the image file size was maximally reduced without visually losing information and inducing blur. Images were then further downsampled to fulfil the image size requirements of QuickNII (scaling factor: 0.50) (detail at: https://quicknii.readthedocs.io/en/latest/imageprepro.html).

#### Image Registration to the CCFv3 with QuickNII and VisuAlign

Serial section images from one brain (irrespective of stain) were combined into a descriptor XML file using the QuickNII Filebuilder application (included in the QuickNII download package). QuickNII (RRID:SCR_016854, QuickNII-ABAMouse-v3-2015 version 2.2) was used to perform linear registration to the CCFv3 2015 followed by nonlinear refinement with VisuAlign (RRID: SCR_017978, version 0.8). For each image series, the thionine-stained sections were registered first since they provided the greatest visualization of region boundaries. Subsequently, all remaining sections were registered in a serial manner. Two independent raters evaluated the registration of each section performed with QuickNII and the refinements made with VisuAlign. Spatial registration data was exported from both QuickNII and VisuAlign in JSON and FLAT files to be used in the Nutil software.

#### Cell Segmentation with ilastik

The ilastik software (RRID:SCR_015246) supports feature extraction by segmentation based on supervised machine learning algorithms. For each stain, ten training images with representative staining were loaded into the Pixel Classification workflow in ilastik (v.1.3.3). Two classes termed “label” and “background” were created, and annotations of each class were applied in all the training images until the segmentation was deemed satisfactory and confirmed by two independent raters. The trained classifiers were applied to all the images of that stain using the batch processing function in ilastik. Segmented images were exported in 8-bit indexed PNG format. Red-green-blue (RGB) colors were applied to the images with the Glasbey Lookup Table in FIJI^48^.

#### Evaluation of Section Image Quality with QCAlign

The QCAlign software (RRID:SCR_023088, version 0.7) was used to assess the integrity of the sections for each brain image series (all 40 brains were assessed) using a 5-voxel grid spacing. This involved marking up points that overlapped areas of damage (representing tears in the tissue, folds, artifacts, and errors in image acquisition) for all sections. Results were exported in TXT format and used to calculate percentage damage per section by dividing the number of damage markers by the total number of markers overlapping the section (damage = # damage markers per section / # of total markers per section). Section images with more than 30% damage were deemed unsuited for QUINT analysis (Supplemental Table 3). Nutil results per brain were re-calculated in R following removal of results from the damaged sections.

#### Creation of a Customized Atlas Hierarchy with QCAlign

Brain reference atlases such as the CCFv3 are organized in systematic hierarchies that group related regions^47^. A customized hierarchy level was created with QCAlign to be used for the quality control assessment of the atlas-registration, and to define customized regions to be quantified (hereafter referred to as the “intermediate hierarchy”). To create this intermediate hierarchy, the atlas delineations supplied by the workflow were overlaid on the thionine-stained sections at the finest level of atlas granularity (full expansion of the CCFv3). A grid of points with a 15-voxel grid spacing was applied to the images, with the registration accuracy of each point marked up based on anatomical expertise (“accurate”, “inaccurate” or “uncertain”). If a region received many “uncertain” markers due to obscure region boundaries, the hierarchy level was adjusted one level up, and the process was repeated until the position of most of the markers could be verified (either “accurate” or “inaccurate”). The customized hierarchy was exported as a TXT file to be used in the Nutil software to define the regions for quantification (Supplemental Table 4).

#### Quality Control Assessment of Atlas-Registration to the Section Images using QCAlign

In the QUINT workflow, 2D atlas-plates are created to match the cutting angle of the sections and registered to the section images in a linear manner using QuickNII. Next, these atlas-registrations are warped (in-plane) to provide a better fit to the sections using VisuAlign. To determine the quality of the atlas-registration to each region in the intermediate hierarchy, ten raters across two academic institutions were recruited to perform a quality assessment using the QCAlign software. Raters varied in anatomical knowledge with expertise ranging from postbaccalaureate researchers, Ph.D. students, senior post-doctoral fellows, and associate research scientists in the field of neuroscience and neuroanatomy. Assessments were performed on the atlas-registration achieved using QuickNII only (2 raters), and on the atlas-registration achieved using both QuickNII and VisuAlign (10 raters). All assessments were performed on the thionine-stained sections from five brains (selected at random) at the intermediate hierarchy level established by the method described above. To perform the assessment, markers with a 15-voxel grid spacing were overlaid on the sections and the position of each marker was assigned as either “accurate”, “inaccurate” or “uncertain” based on anatomical expertise. This was determined by inspecting the position of the marker with respect to visual landmarks in the section and comparing that to the name of the region, which was revealed by hovering over each marker. The atlas-delineations were switched “off” during this assessment because the delineations obscure boundaries in the sections and may bias the outcome.

The QCAlign results were exported in TXT format with counts of accurate, inaccurate, and uncertain markers indicated per region, per section, and per brain. Regional accuracy, inaccuracy, and uncertainty scores were calculated per rater/brain and per brain overall with R-Studio (shared at https://github.com/Neural-Systems-at-UIO/BRAINSPACE). Uncertainty scores were calculated by dividing the number of uncertain markers by the total number of markers in the region, reflecting the percentage of the region for which the registration could not be verified as either accurate or inaccurate (Uncertainty Score = (# uncertain markers)/(# accurate markers + # inaccurate markers + # uncertain markers). Since it was not possible to verify the registration of all the points in the regions (some points were assigned uncertain markers due to a lack of landmarks or limited expertise), the calculation of accuracy and inaccuracy scores correspond to the parts of each region for which the registration could be verified. Thereby, accuracy scores should be inspected together with the uncertainty scores, since a high uncertainty means that the accuracy corresponds to a limited part of the region only. Regional accuracy scores were calculated by dividing the total number of accuracy markers by the total number of accurate and inaccurate markers within that region (uncertain markers did not contribute to this calculation) (Accuracy Score = # accurate markers/ (# accurate markers + # inaccurate markers)). Mean regional accuracy and uncertainty scores were calculated by dividing the summed score of all assessments by the total number of assessments. For each intermediate hierarchy region, the number of assessments contributing to the calculation of the mean accuracy and uncertainty scores depended on the number of raters and number of brains assessed, as well as how often accurate or inaccurate markers could be assigned by the raters (depending on presence of grid markers in that region, tissue quality, and/or anatomical expertise, etc.). In some cases, regions were marked entirely as uncertain by raters; therefore, excluding these assessments from the mean accuracy calculation. For the registration achieved with QuickNII only, a maximum of 10 assessments were averaged across all raters/brains (Brain 1: two raters’ assessments, Brain 2: two raters’ assessments, Brain 3: two raters’ assessments, Brain 4: two raters’ assessments, Brain 5: two raters’ assessments). For the registration achieved with QuickNII and VisuAlign a maximum of 36 assessments were averaged across all raters/brains (Brain 1: ten raters’ assessments, Brain 2: seven raters’ assessments, Brain 3: seven raters’ assessments, Brain 4: six raters’ assessments, Brain 5: six raters’ assessments).

### Regional quantification of stain load with Nutil

Nutil (RRID: SCR_017183) supports regional quantification of IHC-stained features by applying the *Quantifier* feature to combine the output from the atlas-registration (QuickNII and VisuAlign) and feature extraction (ilastik) steps. Nutil (v0.7.0) was used to quantify the percentage of IHC-stained area per region area (hereafter referred to as “load”) in the customized regions defined by the intermediate hierarchy level per stain and brain series. Since hemibrain sections rather than whole brain sections were analyzed in the study, customized masks were created and used to exclude the atlas regions located in the missing hemibrain from the quantification. The hemibrain masks were created with the QNLMask software that is shared with the VisuAlign software (https://www.nitrc.org/projects/visualign). Nutil analysis was performed separately for each stain, with quantification of regional load of neurons (NeuN), microglia (Iba1), reactive astrocytes (GFAP), all nuclei (thionine), and beta-amyloid 1-42 pathology (AB1-42) achieved according to the parameters defined in the NUT file (shared in the BRAINSPACE GitHub repository). The object splitting feature was switched “on” to ensure correct calculation of the regional loads. The NUT files were created and read into Nutil via the command line to batch-process multiple brains in succession. The regional load values obtained from the Nutil reports were used in downstream analysis. Regional load was quantified after QuickNII registration alone, and following QuickNII registration supplemented with VisuAlign refinement. Regional stain loads can either increase or decrease following nonlinear refinement compared to load calculated after QuickNII alone depending on the changes made to regional boundaries, the overall density of pathology or cells in that region, and the stain being evaluated.

### Sample and Region Exclusion from Post Analyses

Data from one female 6m mouse of AD-BXD strain 44 was removed from the downstream analysis because the majority of the sections were severely ripped prohibiting successful atlas-registration. Quantification output from all of the 77 regions in the intermediate hierarchy file are included in the Nutil reports (shared as the BRAINSPACE project on EBRAINS Knowledge Graph Search, https://search.kg.ebrains.eu/). In the present study, 55 of these regions were included in the QCAlign assessment of the atlas-registration across 5 brains; and 43 of these regions were included in the assessment of cell and pathology load across 37 brains (5XFAD mice only). Specific region exclusion criteria are reported in Supplemental Table 5. As a brief summary, some of the atlas regions did not have results in the reports since they were not represented in the sections or corresponded to a parent structure with results provided at a finer level of atlas granularity. Regions with no biological results were disregarded from all analyses. Furthermore, results from several regions were not analyzed in the present study due to low representation in the sections.

### Statistical analysis of QUINT data

For each stain, the load values of 43 intermediate hierarchy provided by the Nutil software were used for comparative analysis across 5XFAD brains at 6 m (n = 17) and 14 m (n = 20). Data have been expressed as means ± standard error of the mean (SEM) or as otherwise indicated in graphs. Statistical analysis of data was performed using R version 4.0.0 (2020-04-24) -- “Arbor Day”. Wilcoxon two-way assessment (strain and age factors) was implemented to determine if there were significant differences in the stain load as registered using QuickNII alone vs registered using QuickNII and VisuAlign. Analysis of variance (ANOVA) (age and strain factors) was used to determine whether there were significant differences in regional stain load between 6m and 14m groups. Multilevel Pearson correlations with and without age corrections were used to evaluate the relationship between hippocampal stain load and gene expression. Multiple testing corrections for each test was performed using false discovery rate (FDR) correction. Criterion for measures to be considered uncorrected significant was p-value < 0.05 and significant after correction was FDR p-value < 0.05.

### Immunohistochemistry and Bulk RNA Sequencing Integration

To identify genes associated with variation in hippocampal cell and pathology load we integrated our IHC quantification with RNAseq data. The goal of this analysis was to determine whether changes in cell composition contributed to subsequent changes in hippocampal gene expression detected via RNAseq. Only 5XFAD samples with paired IHC and RNAseq data were selected (n =34); therefore, all animals in this analysis had one hemisphere fixed for IHC and the contralateral hippocampus dissected for bulk RNAseq. The RNAseq data used in the current study was previously published and the dataset series (GSE) are accessible via the National Center for Biotechnology Information Gene Expression Omnibus (GEO) (GEO: GSE101144, GEO:GSE119215, GEO:GSE119408)^11, 13, 16^. Expected read counts (ERCs) were filtered to include genes with >10 ERCs in more than 50% of the samples from 5XFAD mice, resulting in 15,703 of 47,645 genes that passed filtering. Following the exclusion of genes with low read counts, datasets were batch-corrected using the R Combat-Seq package, then normalized and transformed using the default pipeline of R DESeq2^49^. The relationship between gene expression and stain load (AB1-42, NeuN, GFAP, and Iba1) from the hippocampal formation summary region was assessed using Pearson’s correlation from linear mixed models^50^, which allowed the effect of age on the association between gene expression and load to be accounted for by including age as a random effect (correlation(partial = TRUE, multilevel = TRUE). P-values per stain and gene correlation were corrected for multiple comparisons via FDR correction and considered significant if the FDR p-value < 0.05. Genes that were exclusively significantly correlated (uncorrected p-value < 0.05) prior to age adjustment were deemed to be age-dependent correlates. Genes that were exclusively significantly correlated (uncorrected p-value < 0.05) following age adjustment were deemed to be age-independent correlates. Gene Set Enrichment Analysis (GSEA) queried against Reactome pathways was carried out in WebGestalt^51–54^ using the output correlation coefficients per gene and stain for each multi-level correlation method (age-adjusted and non-age-adjusted). Advanced GSEA parameters used included: Minimum number of IDs in the category: 20, Maximum number of IDs in the category: 2000, Significance Level: FDR < 0.05, and Number of permutations: 1000). Lastly, individual ERC and hippocampal load data were incorporated into a DESeq model, and the design was run on the intercept (∼1). Transformed normalized counts for boxplots in figure 5 were obtained using the DESeqDataSetFromMatrix() and counts() functions. Scripts used for RNAseq normalization and modeling, IHC and RNAseq correlations and visualization can be accessed on GitHub at:https://github.com/Neural-Systems-at-UIO/BRAINSPACE/tree/main/Scripts.

### Data Availability

The collection of section images, accompanying meta data, atlas-registration files and output, as well as Nutil output are shared as the BRAINSPACE project via the EBRAINS Knowledge Graph Search (https://search.kg.ebrains.eu). R scripts used to complete statistical analyses are publicly available on GitHub at: https://github.com/Neural-Systems-at-UIO/BRAINSPACE.

### Sharing of QUINT tools and disclaimer

All the software in the QUINT workflow are open-source and shared on GitHub and nitrc.org under MIT license for QuickNII and VisuAlign; GNU General Public License (GPL) v3.0 for Nutil; and GPL v2 / GPL v3 for ilastik. While the software are validated based on multiple ground truth datasets shared on the Nutil GitHub page, we recommend independent validation of data from QUINT prior to use. To validate the QUINT workflow for the present study, Nutil v0.7.0 was used to analyze two synthetic datasets with objects of known size and anatomical location based on the parameters selected for the study. The validator feature in Nutil confirmed that the results were identical to the ground truth. The dataset, ground truth and results of Nutil v0.7.0 are shared on GitHub at https://github.com/Neural-Systems-at-UIO/BRAINSPACE/tree/main/Nutil_Validation. The QUINT workflow is shared on EBRAINS (<u>ebrains.eu/service/quint</u>), with user documentation (https://quint-workflow.readthedocs.io) and user support available through EBRAINS.

## Results

### New functionality added to the QUINT workflow supports high-throughput analysis of diverse AD-BXD strains

The original QUINT workflow was designed to support the quantification of IHC-stained features in images of serial brain sections by linear registration to a reference brain atlas in combination with feature extraction by supervised machine learning^38^. While this method works well for serial sections that closely resemble 2D atlas-planes throughout the reference atlas template (typically generated based on intact whole brain tissue); in practice, the technical procedures of fixing, sectioning, staining, and mounting sections often lead to distortions, tears in the sections, and artifacts that impact the quality of the linear atlas-registration. Since reference atlases are created based on standard reference animals (young adult male B6 mice in the case of the CCFv3)^47^, sections originating from strains and/or ages that genetically differ from such animals may also have anatomical differences relative to the reference template. Recognizing the need to customize the linear atlas-registration and provide a better match of the atlas overlay on individual sections, a new tool that supports nonlinear refinement was created and incorporated in the workflow (VisuAlign) (Figure 1a). Nonlinear refinements are manually applied based on visual landmarks in the sections. Furthermore, a quality control tool based on systematic random sampling was created for validating the quality of the atlas-registration to each region (QCAlign). This manual assessment is based on the overlap between the delineations supplied by the atlas and landmarks revealed by IHC staining. Since only a limited number of landmarks can be revealed by IHC staining, a method was also implemented for adjusting the granularity of the reference atlas to a level that supports the verification of the atlas registration. This functionality of QCAlign provides users a platform for flexible assessment of the Allen Mouse Brain Atlas, which can be manipulated to display a complete or reduced atlas hierarchy overlaid on the sections. Individual reference atlas regions can be compiled into larger themed regions (e.g. isocortex), allowing users to tailor the assessment to their unique experimental design and research interests. Lastly, since there are other factors that can affect the quality of the results that can be achieved with QUINT (for example, artifacts that obscure the staining, or tissue damage too extensive to account for by nonlinear warping), a method within QCAlign was also introduced to promote the systematic screening of sections, and for assessing their suitability for QUINT analysis. This feature is particularly useful in the context of high-throughput studies since it allows exclusion of sections according to systematic criteria. The expanded QUINT workflow was applied to serial section images from the diverse AD-BXD mice (Figure 1b) to quantify all nuclei (thionine), neurons (NeuN), microglia (Iba1), reactive astrocytes (GFAP) and amyloid beta pathology (AB1-42) in customized regions compiled from CCFv3 regions. Examples of these IHC-stained sections are shown in Figure 1c. Each step of the QUINT workflow generates a visual output that can be shared together with the final results of the workflow to support independent verification of findings (Examples of the visual output are shown in Figure 1d).

### Quality of the atlas-registration performed in the QUINT workflow can be confirmed using QCAlign

The new QCAlign tool was implemented to assess the quality of the atlas-registration achieved using QuickNII and VisuAlign. First, the full CCFv3 2015 was condensed into 77 regions to create an intermediate hierarchy of regions that was exported from the QCAlign software (Supplemental Table 4). These regions have visually discernable boundaries as detected in the thionine-stained sections (example images with superimposed atlas-delineations are shown in Supplemental Figure 1a). Next, with the hierarchy level set in QCAlign, a rater can perform an independent assessment and rate the accuracy of the atlas-registration as performed in the workflow (Supplemental Figure 1b). This entails assigning grid markers positioned at a set density over the sections as either accurate, inaccurate, or uncertain based on anatomical expertise (Figure 2a). A grid point is marked as “accurate” if the assigned atlas-registration correctly matches the region depicted in the section. This is determined by the investigator based on landmarks; therefore, the region boundaries in question must be distinct enough to make this call. If there is a discrepancy between the registered atlas region and what the rater identifies the region to be in the brain section, the “inaccurate” marker is assigned. Inaccurate markers can be the result of incorrect registration using QuickNII, and/or incomplete adjustment during VisuAlign refinement. If a high frequency of inaccurate markers is assigned, the initial registration of brain sections should be reevaluated. Lastly, an “uncertain” marker is placed when the rater lacks the anatomical knowledge to apply an accurate or inaccurate marker with confidence, or when the borders between regions are ambiguous hindering the ability to differentiate regions. If a high frequency of uncertain markers is assigned, the rater should reconsider the hierarchy level chosen for the evaluation.

**Figure 2.**
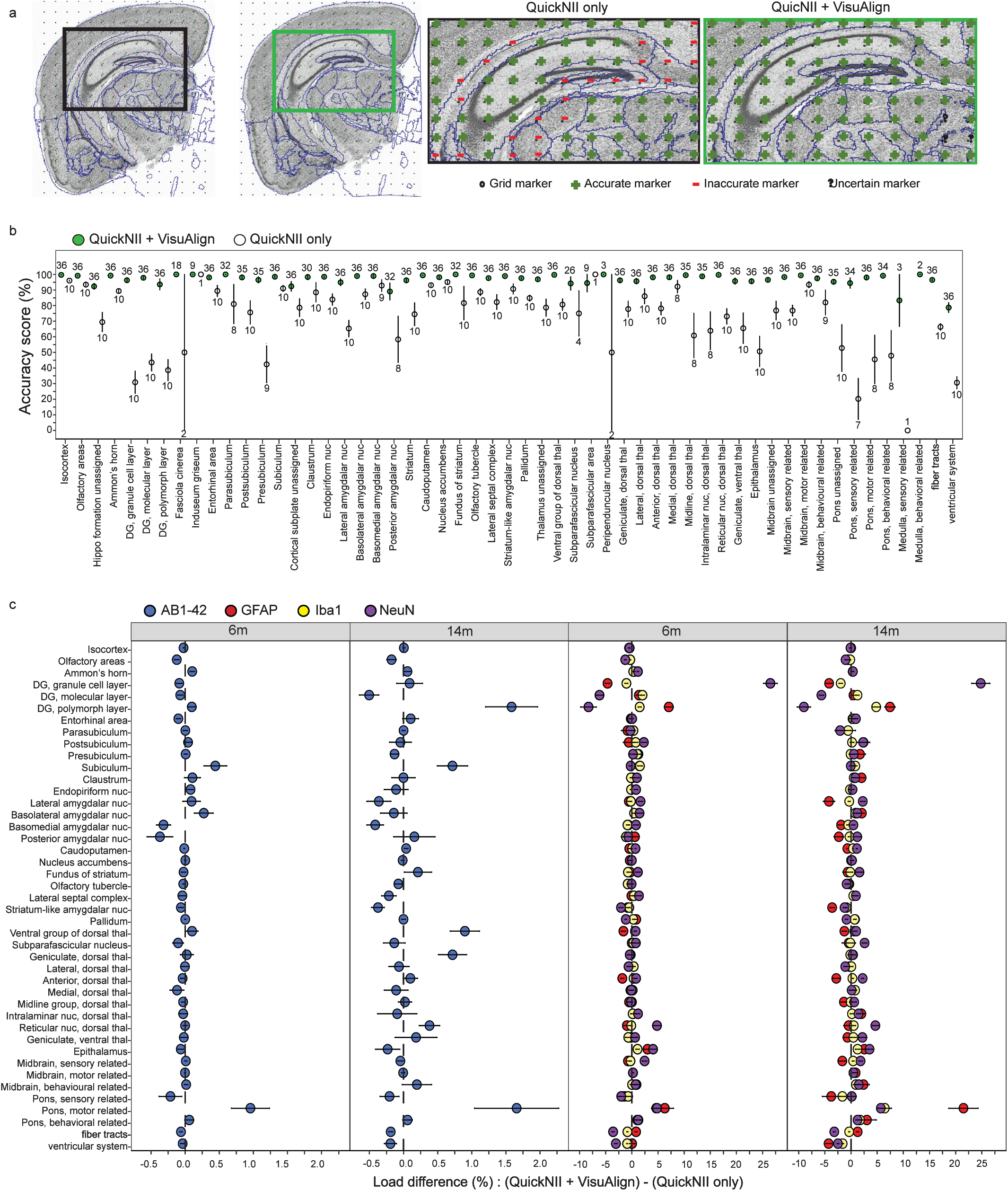
QCAlign verification of regional atlas-registration at the selected intermediate hierarchy level **a.)** QCAlign quality control assessment can be completed after rigid QuickNII registration alone or following the use of QuickNII and VisuAlign to verify the registration to each region in the sections. Inset) Example of completed QCAlign assessment in the hippocampal formation after QuickNII Only and QuickNII + VisuAlign registration. **b.)** Mean accuracy scores per intermediate hierarchy region after QuickNII registration alone (white) or after QuickNII and VisuAlign registration (green). Two raters scored the same 5 randomly selected brains after QuickNII registration alone, max n=10 per region (Raters: n= 2 per brain). Up to 10 raters scored the same 5 randomly selected brains after QuickNII and VisuAlign registration, max n=36 per region (Raters: n= 6-10 per brain). Dots represent the mean score across raters per region for 5 brains ± SEM, with the numbers labels representing the number of assessments contributing to each calculation (QuickNII alone labels are below white points, QuickNII + VisuAlign labels are above green points). **c.)** The impact of VisuAlign refinement on regional stain load (%-stain-positive coverage/per region area) was measuring by calculating the difference in load following Nutil quantification after each method (regional (QuickNII + VisuAlign output (%) – regional (QuickNII output(%) = regional load difference (%)). Dots represent mean regional load difference ± SEM for all 5XFAD animals at 6m and 14m (6m: n=17, 14m: n=20).

To confirm the atlas-registration following VisuAlign adjustment, ten researchers across two academic institutions were recruited to perform a quality control assessment of atlas-registrations using QCAlign. The assessment was performed on the thionine-stained sections from 5 brains selected at random from the cohort of 39 brains. A maximum of 36 assessments were averaged per intermediate hierarchy region (6-10 raters assessing up to 5 brains) (Supplemental Figure 2). There was high consensus among raters that the registration to the intermediate hierarchy regions was highly accurate (100%-78.7% accuracy score) (Figure 2b, green). Regions with the greatest accuracy scores were regions compiled of many subregions (e.g. isocortex, 99.7%, SEM ± 0.057) and/or that have very distinct anatomical borders (e.g. caudoputamen, 99.4%, SEM ± 0.129). Smaller regions had the potential to have zero grid markers randomly placed within their area resulted in reduced number of assessments contributing to the mean accuracy score (e.g. subparafascicular area, n= 9 assessments). Regions with the lowest rater sampling rate were among the regions with the highest variation and lowest accuracy scores. Regions with appropriate rater sampling (n>20 assessments) but low accuracy scores included the posterior amygdalar nucleus (89.1%, SEM ± 5.37) and the ventricular systems (78.7%, SEM ± 3.11). The low accuracy attributed to the posterior amygdalar nucleus could be due to its relatively ambiguous border with the posterior olfactory area and the subiculum. Also, regions of the ventricular system were consistently difficult to align in both QuickNII and VisuAlign since they are prone to distortion (e.g. lateral ventricle) or are located in medial locations along the midline where the brain was bisected into hemibrains (e.g. third ventricle), resulting in low accuracy overall. To summarize, we created a new tool for quality control assessment of the atlas-registration and, by using this tool, were able to confirm the ability of the QUINT workflow to achieve highly accurate registration of the regions in the intermediate hierarchy.

### Nonlinear adjustment increases regional registration accuracy, and impacts cell and pathology load estimates

VisuAlign offers the unique ability to refine and improve the atlas-registration to diverse AD model mouse brain sections by allowing users to make nonlinear adjustments to the atlas plates set in QuickNII. The importance of completing nonlinear warping following linear registration was highlighted by comparing the QCAlign output following each atlas-registration step in the QUINT workflow (Figure 2a). Linear registration achieved using QuickNII alone is susceptible to error as indicated by the higher frequency of inaccurate markers. The hippocampus is a particularly vulnerable region that requires non-linear adjustment due to the distinct shape and relatively small size of the dentate gyrus (Figure 2a inset). Regional accuracy scores of five brains were calculated and compared following atlas-registration performed using QuickNII only (2 raters) relative to the registration performed using QuickNII then adjusted in VisuAlign (6-10 raters) (Figure 2b). The completion of nonlinear warping in VisuAlign greatly improved the registration of atlas regions to the brain sections (Figure 2b, green vs white). Regions that exhibited the greatest increases in accuracy scores included those that are often not prioritized when initially aligning atlas plates to the brain sections in QuickNII, thereby requiring more extensive nonlinear adjustment (i.e. regions comprising the mid-and hindbrain). Regional quantification of cellular and pathology load was also impacted by the increased accuracy of registration achieved following nonlinear warping. Regions that required the most adjustment in VisuAlign, thereby exhibiting the greatest increases in accuracy, also had the greatest difference in load values when comparing regional load output from registration using QuickNII alone versus registration completed in QuickNII and refined in VisuAlign (Figure 2c, Supplemental Table 6).

### AD-BXD strains exhibit widespread increases of glial and amyloid pathology from 6m to 14m

Differences in cell composition and amyloid pathology load were compared between 5XFAD carriers of 6m and 14m to detect regional changes that occur with age and AD (Figure 3, Supplemental Table 7). Among 5XFADs, there are only minor changes in NeuN load between 6m and 14m animals overall (Figure 3, i). The only regions that exhibited significant age-related (FDR-corrected p-value<0.05) decreases in NeuN load were the Ammon’s horn(p-value=0.0472) and dentate gyrus, polymorph layer(p-value=0.00299). Slight, but significant (FDR-corrected p-value<0.05) increases in NeuN load were observed with age in the posterior amygdalar nucleus (p-value = 0.0327) and striatum-like amygdalar nuclei (p-value=0.0258) (Figure 3a, i). Increased glial proliferation and reactivity are also hallmark symptoms of AD progression with age. Within this dataset, we confirmed that regional astrocyte and microglial cell load increased from 6m to 14m in 5XFAD animals. Regionally, the caudoputamen exhibited the most significant increases in GFAP load (p= 2.91E-10, FDR-corrected) (Figure 3a, ii). The midbrain (motor-related) regions (FDR-corrected p-value= 1.26E-08) and olfactory tubercle (FDR-corrected p-value=1.55E-08) exhibited the greatest microglial load increase from 6 to 14m (Figure 3a, iii). Aligned with previous reports in 5XFAD animals, amyloid pathology was most prevalent within the subiculum at the earlier 6m time point^55^ (3.41% ± 0.227% SEM, Figure 3a, iv). In addition to the subiculum, amygdalar regions were highly susceptible to increased amyloid deposition by adulthood (6m) (Figure 3a, iv). As an aggressive amyloidosis AD model, the 5XFAD animals exhibited a near global increase in amyloid deposition between 6m and 14m. Amyloid deposition was strongly associated with the hippocampus and hippocampal-projected regions, including the cortex, thalamus, and amygdalar regions as previously noted (Figure 3a, iv)^56^. All regions besides the claustrum, lateral amygdalar nucleus, parasubiculum, midbrain (behavioral state related), pons (behavioral state related), pons (motor related), and pons (sensory related) regions exhibited a significant increase in amyloid load from 6m to 14m.

**Figure 3.**
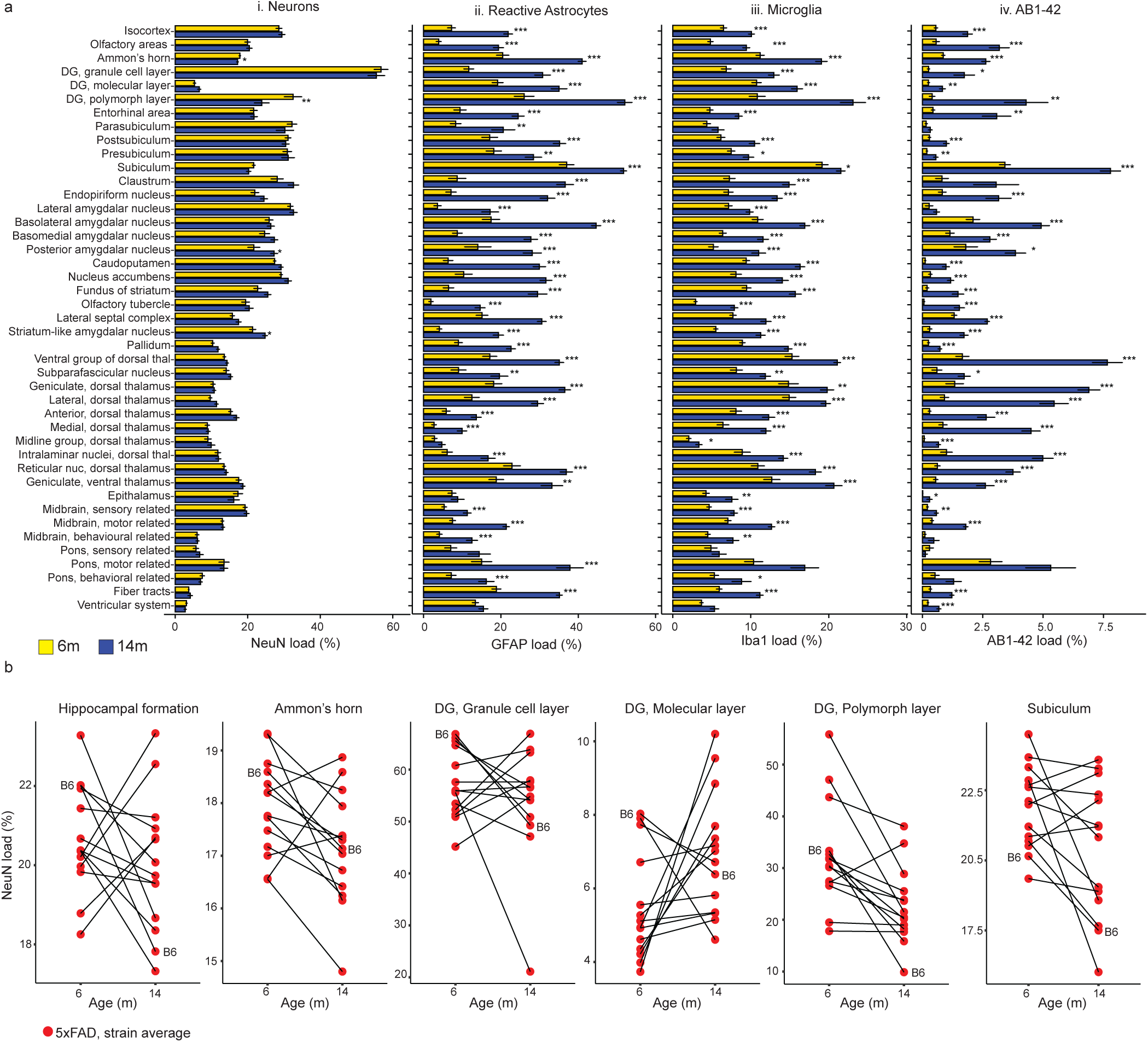
Regional pathology and cell load vary from adulthood (6m) to middle age (14m) in 5XFAD mice. **a.)** Regional cell and pathology load of the intermediate hierarchy regions of 5XFAD mice. i. Differences in NeuN load between the age groups were limited across the intermediate hierarchy regions. ii-iv. GFAP, Iba1, and AB1-42 load increased with age across most intermediate hierarchy regions. Bars represent regional averages ± SEM for 6m and 14m groups. (5XFAD mice only, 6m: n=17, 14m: n=20). FDR corrected p-values represented. P-value: * <0.05, ** <0.01, *** <0.001. **b.)** Strain averages of NeuN load across the hippocampal formation and hippocampal intermediate hierarchy subregions. Points are mean load per strain. Lines connect strain matches across the two age groups: 6m and 14m. Only strains with an aged match counterpart are represented (5XFAD mice only, 6m: n=17, 14m: n=18, n= 1-3 per strain). The B6 founder strain is labeled for reference.

### Individual AD-BXD strains exhibit variation in region neuronal load

The hippocampus, known as a structure involved in cognitive processing, memory formation and storage^57^, has been elaborately studied in the context of aging and AD. Compared to the near global increase in glia and pathology among AD-BXD strains between 6m and 14m, fewer age-related differences in neuron load were detected (Fig 3a, i). Of the four regions that displayed a significant difference in NeuN load between 6m and 14m after FDR correction, two of those regions were within the hippocampus. While regional variation in NeuN load was minimal overall within the age groups, age-related strain-specific variation was revealed by investigating changes in NeuN load in hippocampal subregions on a per strain basis (Figure 3b, Supplemental figure 3). AD-BXD strains displayed a range from neurodegeneration to neuronal maintenance between 6m and 14m, modeling the heterogeneity observed in human AD^58^. No strain effect was detected in stain load among the 43 intermediate atlas regions quantified (uncorrected p-value> 0.05, 2-way ANOVA), but since sample sizes per strain were relatively small in this analysis, a potential strain effect cannot be firmly excluded and will be evaluated when the sample size is increased in future analyses.

### Integration of paired IHC and bulk RNA sequencing data reveals cell load is a confounding factor in age by gene expression correlations among AD-BXDs

Using the QUINT workflow, we reported variation in cell and pathology load between age groups and among AD-BXD strains (Figure 3). As mentioned earlier, due to the inherent properties of bulk RNAseq, which allow for single, tissue-averaged, gene expression measurements, the influence of cell composition is often overlooked in the interpretation of analyses and may conflate expression differences driven by other experimental factors such as age and pathology^27, 59, 60^. Here, using output from our QUINT workflow analysis, we demonstrate that ∼15-35% of genes expressed in the hippocampus are correlated with load and vary based on both cell-type and age. To do this, we integrated hippocampal formation cell (NeuN, GFAP, Iba1) and pathology (AB1-42) load output with gene expression data measured via bulk RNAseq obtained from the contralateral hippocampus of the same mice at two age time points (previously published^11, 13, 16^).

Hippocampal load per stain type (NeuN, GFAP, Iba1, and AB1-42) was correlated with normalized read counts to identify age-dependent relationships between load and gene expression. The percentage of the 15,703 genes analyzed in the RNAseq dataset that were significantly correlated (uncorrected p-value < 0.05) with load varied by stain type (NeuN: 16.35%, GFAP: 36.76%, Iba1: 34.78%, AB1-42: 31.86%) (Figure 4a; labeled genes indicate the top 5 positively and 5 negatively correlated significant genes (FDR-corrected p-value < 0.05). Non-coincidentally, stains that had the most significant gene correlates had the greatest age-related changes in load. Since our population is comprised of mixed ages and age is a primary driver of variation in load (Figure 3), this effect of age may be masking genes that are related to load in an age-independent manner. We aimed to elucidate this subset of genes by testing the role of age as a mediator of the relationship between stain load and gene expression in our 5XFAD population by using a multilevel correlation approach adjusting for the effect of age. Similar to the outcomes of the age-dependent correlation above (Figure 4a), the percentage of genes significantly correlated after age adjustment (uncorrected p-value < 0.05) with load also varied by stain type (NeuN: 12.56%, GFAP: 23.53%, Iba1: 18.30%, AB1-42: 12.34%, Figure 4b). The number of correlated genes (uncorrected p-value < 0.05) was reduced following age-adjustment across all stains, with AB1-42 exhibiting the greatest reduction of significantly correlated genes (19.52%, Figure 4a, 4b). Next, we sought to differentiate genes that were exclusively correlated with load either before or after age-adjustment. By further comparing both analyses (age-unadjusted, Figure 4a and age-adjusted, Figure 4b), we classified genes into 1) exclusively significantly associated with variation in load in an age-dependent manner (non-age-adjusted output (orange in figure 4c)), 2) exclusively significantly associated with load irrespective of age (age-adjusted output (blue in figure 4c)), or 3) significantly associated with both load and age (non-age-adjusted and age-adjusted output (green in figure 4c)) (Supplemental Table 8). The majority of correlations between gene expression and load were driven by age as indicated by the greater abundance of non-adjusted significant genes per stain (Figure 4c). This age-driven relationship is illustrated by the correlation between Iba1 load and polypeptide N-acetylgalactosaminyltransferase 6 (*Galnt6*) expression, which was identified to be a top gene that is highly associated with variation in Iba1 load in an age-dependent manner (Figure 4d, i). *Galnt6* has been found to have increased mRNA expression in the brains of AD patients and be related to AB production^61, 62^. Here, *Galnt6* exhibited increased expression with age that parallels the increase in Iba1 load observed from 6m to 14m (Figure 4d, i-ii). This trend of increased load matched by a change in gene expression between 6m and 14m was unique to the most highly correlated genes prior to age adjustment. On the contrary, 0.78%-5.86% of the genes per stain were exclusively significant only after age-adjustment, indicating that these genes are likely associated with load in an age-independent manner (Figure 4c). These age-independent genes exhibited a pattern of increased cell (GFAP and Iba1) and pathology (AB1-42) load but no difference in gene expression between 6m and 14m. This pattern is exemplified by looking at the relationship between gene expression and load with age for transmembrane protein 39A (*Tmem39a*), a topmost correlated gene with Iba1 load after age-adjustment (Figure 4d, ii). *Tmem39a* is a known contributor to pathways implicated in AD, including inflammation, dysregulated type I interferon responses, and other immune processes^63^; and like other highly correlated genes following age-adjustment, *Tmem39a* exhibited specific within-age-group associations between load and gene expression (Fig4d, ii). These genes with stronger significance following age adjustment may be driven by load differences seen between the groups independent of the effect of age on load. Identifying and differentiating age-dependent and age-independent gene correlates promotes the prioritization of gene candidates and recognition of whether the expression of these genes are relative to the proportions of different cell types that are altered with age and AD.

**Figure 4.**
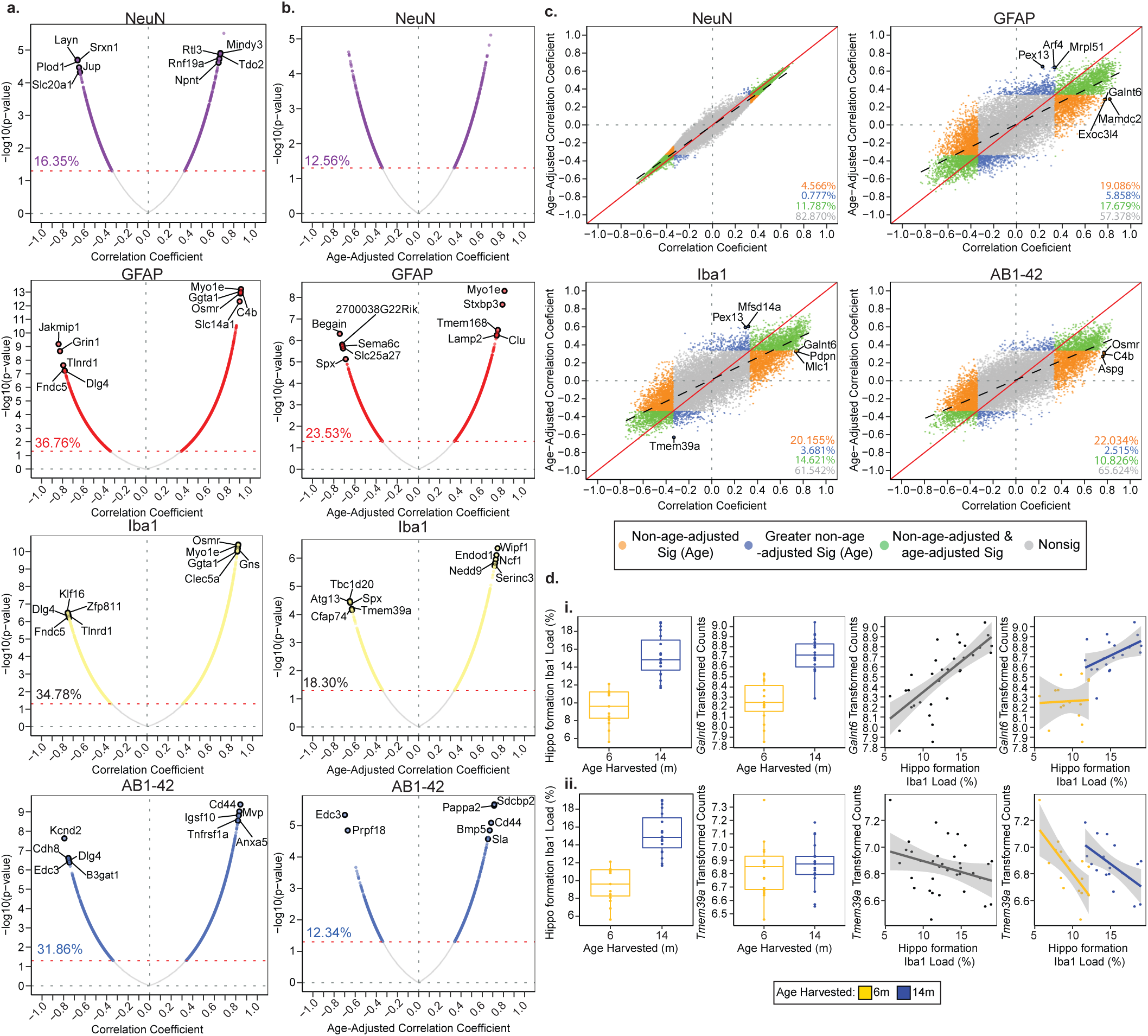
Stain-specific load correlations with RNAseq gene expression to identify genes impacted by changes in load within the hippocampal formation. **a.)** Gene expression by load Pearson R correlation coefficients and p-value relationships without age adjustment for each stain. Significantly correlated genes (uncorrected p-value < 0.05) are colored in each plot. The percentage of uncorrected significant genes is indicated within the plot. The top five positive and negative FDR significant (FDR p-value < 0.05) correlated genes are labeled. **b.)** Gene expression by load Pearson R correlation coefficients and p-value relationships after age adjustment for each stain. Significantly correlated genes (uncorrected p-value < 0.05) are colored according to stain. The percentage of uncorrected significant genes is indicated within the plot. The top five positive and negative FDR significant (FDR p-value < 0.05) correlated genes are labeled. **c.)** Comparison of Pearson R correlation coefficients without and with age adjustment per stain. Gene correlations that were exclusively significant (uncorrect-p-value < 0.05) without age adjustment are considered age-dependent (orange). Gene correlations that were exclusively significant (uncorrect-p-value < 0.05) with age adjustment are considered age-independent (blue). The specific influence of age and load cannot be disseminated in gene correlations that were significant (uncorrect-p-value < 0.05) under both correlation conditions (green). All nonsignificant (uncorrect-p-value < 0.05) genes are labeled in gray. The percentage of significant genes per category is represented in the bottom right corner. The top 3 most significant genes per correlation method category are labeled per stain plot (FDR p-value< 0.05). **d.)** Individual relationship between gene expression and load with age for the top age-dependent and independently correlated genes with Iba1. i. *Galnt6* was exclusively significantly correlated with Iba1 without age adjustment. An increase in Iba1 load and *Galnt6* expression occurs between 6m and 14m. A positive relationship between Iba1 load and *Galnt6* expression exists across both age groups as well as within each age group. ii. *Tmem39a* was exclusively significantly correlated with Iba1 after age adjustment. An increase in Iba1 load but not in *Tmem39a* expression occurs between 6m and 14m. A weak relationship between Iba1 load and *Tmem39a* expression exists across both age groups, but separate age-specific correlations with load and gene expression exist. 5XFAD mice only, 6m: n=17, 14m: n=20.

### Mediation of age reveals differential overrepresentation of Reactome pathways

Next, using the correlation coefficients displayed in Figure 4a and b, gene set enrichment analysis (GSEA) was performed to identify pathways that may be biased by individual differences in cell and pathology load (Figure 5). As expected, immune pathways were highly enriched for GFAP, Iba1, and AB1-42 correlations. We also observed a negative relationship between the enrichment of neuronal pathways and GFAP, Iba1, and AB1-42, highlighting the potentially detrimental impact these cell types may have on neuronal functioning in the context of AD. Fewer significantly enriched pathways were associated with NeuN load (age-adjusted and non-age-adjusted), consistent with the subtle changes in load between 5XFADs of 6m and 14m. The most highly enriched pathways for each stain and method (as labeled on the right of the heatmap) were involved in chromatin organization, extracellular matrix organization, immune system, metabolism of RNA, and the neuronal system. In comparing enriched pathways for age-adjusted and non-age adjusted correlations per stain, the greatest difference in the presence of significantly enriched pathways was observed within the cell cycle category for Iba1, GFAP, and AB1-42 stain types. The enrichment of these pathways is consistent with the proliferation of these cell types and pathology and the potential increase in immunoreactive cell cycle proteins^64^. A total of 42 cell cycle pathways were represented across these stain types after age adjustment, while only 2 are present prior to adjustment. Moreover, many negatively enriched pathways including those in the gene expression (transcription) and metabolism of RNA parent pathways were observed almost exclusively within the non-age-adjusted category for GFAP, Iba1, and AB1-42. This pattern of enrichment suggest a more pronounced involvement of these types of pathways with AD-related deterioration with age than necessarily with increased glial and pathology composition^65, 66^. Ultimately, by using these methods we have begun to disseminate the effects of cell and pathology composition in the hippocampal formation and their implication in biologically relevant pathways.

**Figure 5.**
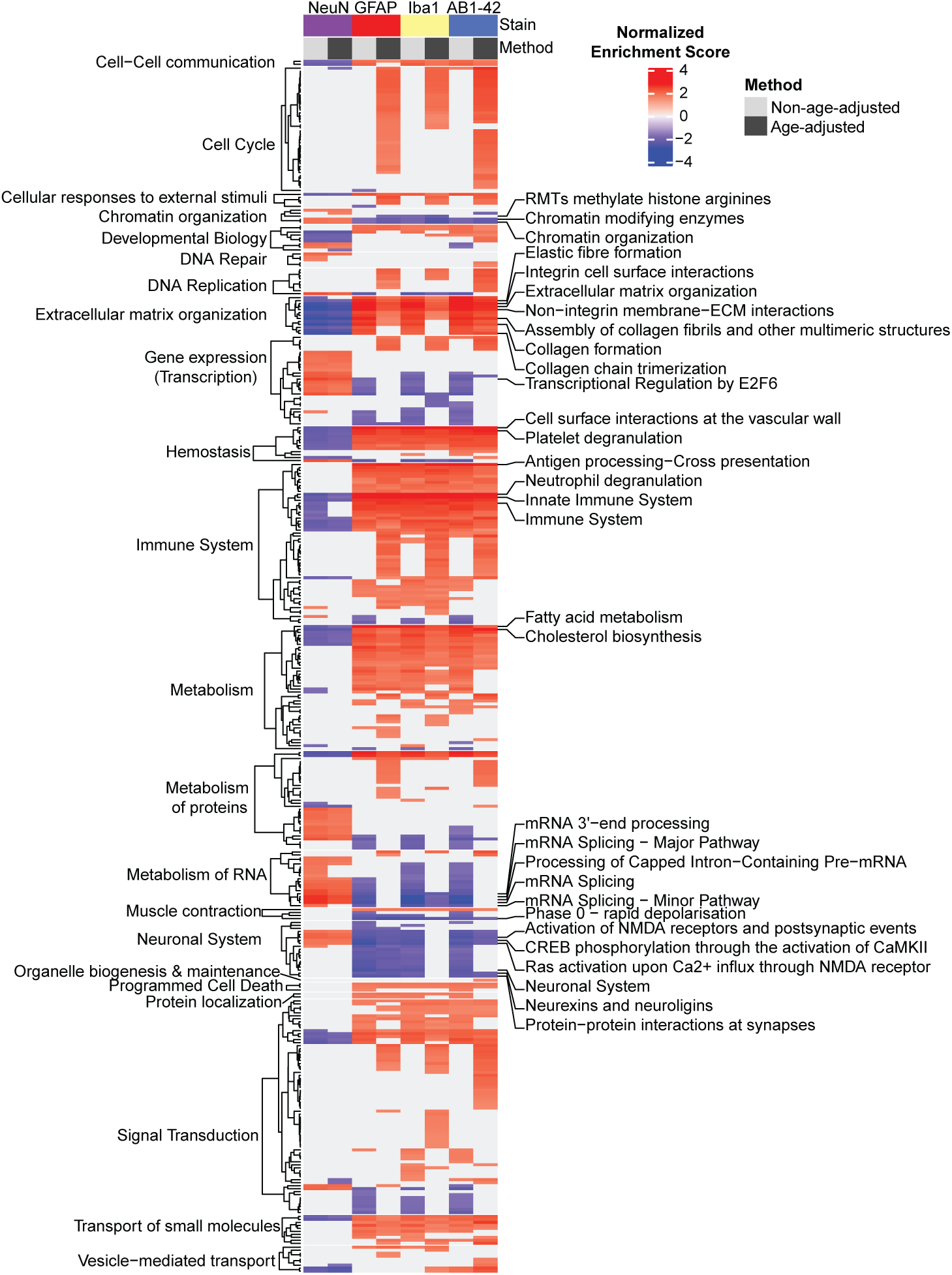
Gene Set Enrichment Analysis (GSEA) of gene correlations per method categorized by Reactome parent pathway. **a.)** Pearson R correlation coefficients from Figure 4a and Figure 4b were input into WebGestalt GSEA to obtain significantly enriched pathways associated with each stain and correlation method (normalized enrichment, non-age-adjusted and age-adjusted). The top three most significant pathways per stain and methods are labeled (FDR p-value< 0.05) (right).

## Discussion

Here, we report on the output from IHC sections of 37 mice from the AD BXD-panel obtained using the expanded QUINT workflow. By adding new functionality to the QUINT workflow to enhance the atlas-registration and perform quality control assessments, we increased the quality of the regional quantification^11^. We quantified age-related differences and characterized the influence of genetic diversity among AD-BXD strains on NeuN, GFAP, Iba1, and AB1-42 load across a validated list of Allen Mouse Brain Atlas CCFv3 2015 subregions^44^. The importance of recognizing this variation in cell and pathology composition was also reflected when integrating gene expression and cell composition data from varying AD-BXD strains. The mouse panel used in this study is considered translationally relevant since it includes strains that incorporate high risk AD mutations (5XFAD) on backgrounds of genetic diversity, thus better recapitulating the complex genotype-phenotype interactions in humans that contribute to symptom variability. The AD-BXD panel provides a unique platform for exploring the effect of genetic background variability on resilience to neurodegeneration, gliosis, and pathology with the potential to reveal resilience genes or pathways that could be targeted for therapeutics.

We demonstrate the capacity of the QUINT workflow to effectively detect subtle differences in regional loads in an accurate manner across the whole brain, which is paramount in the context of high-throughput imaging studies that incorporate genetic diversity models of disease. The quantification of these brains was made possible through the expansion of the QUINT workflow through the development of VisuAlign and QCAlign, as well as through the addition of new functionality to the existing Nutil software. The VisuAlign and QCAlign software were added to the QUINT workflow for a number of reasons. While linear atlas-registration is a useful first step, it often does not produce the required registration precision^67–70^. VisuAlign provides the capability to make nonlinear adjustments to the linear atlas-registration achieved using QuickNII, thus correcting for distortions in the sections introduced during the IHC section preparation as well as for structural differences among brain regions in diverse disease models and age groups. The importance of applying nonlinear refinements was demonstrated by the regional differences in accuracy scores and loads achieved with QUINT-based registration using QuickNII only, relative to registration using QuickNII and VisuAlign. Moreover, since changes driven by genetic differences across strains are likely to be subtle and region-specific, it was crucial to have a method for verifying that the atlas-registration output was accurate. This verification was provided by the QCAlign tool. The limited variability in QCAlign accuracy scores between raters and brains quantified in our 5-brain assessment heighten our confidence that the present cohort of brains was consistently registered to a high standard. Another key functionality of QCAlign is its ability to produce customized hierarchies, which aid in compensating for the difficulty of accurately registering small regions that lack anatomical boundaries. To combat this issue, many investigators generate lists of regions of interest (ROIs) that consist of compiled subregions^71–73^. Our QCAlign tool offers the functionality to create these customized hierarchies by parsing through the 461 regions of the CCFv3 2015 and selecting subregions to compile into related summary regions. Creating a custom hierarchy file from the standard atlas in QCAlign also promotes the labeling of consistent ROIs among laboratories and the ability for anatomists to subsequently verify that the regions selected in the chosen hierarchy are correctly aligned during the registration process. The final feature of QCAlign was developed to detect sections or regions not suited for QUINT analysis due to damage (tears, folds, etc.), artifacts, errors in image acquisition, or other reasons that could potentially skew results. Percentage damage per section or per region can be calculated after marking up sections in QCAlign with the damage marker. This calculation supports removal of results according to transparent, systematic, and reproducible criteria for streamlined high-throughput application.

Overall, the QUINT workflow has a number of advantages over alternative methods. Utilization of the QUINT workflow promotes comprehensive regional analysis as defined by a standardized reference atlas, which facilitates comparison, integration, and reproducibility of results across studies in compliance with the FAIR guiding principles^36, 37^. The ability to share the intermediate results of the workflow (the atlas maps and segmentations) as well as the final results (Nutil Quantifier output) provides transparency and open science, which is important since the atlas-registration and feature extraction steps are inherently subjective processes guided by user-based expertise. Traditional IHC analysis methods that rely on manual delineation of brain regions and counting via stereology are inefficient for brain-wide exploration in studies with large numbers of animals^74–77^. As demonstrated in the present study, the QUINT workflow has the capacity to characterize transgenic models of disease of varying strains, ages, and genotypes; and is designed to support large-scale comparative studies^78, 79^. The workflow is customizable, enabling analysis at different levels of atlas granularity, and with optional features such as the application of masks for hemisphere or other region-based comparisons. Also of note, QUINT is highly accessible irrespective of coding ability since all the steps are performed in software that have user-friendly graphical user interfaces (GUIs). While the subjective nature of the registrations tools is a limitation of the QUINT workflow, it is countered by the addition of the QCAlign software that provides a means to evaluate and document the quality of the atlas-registration performed in the workflow. Another limitation of QUINT is that nonlinear VisuAlign adjustment can be labor intensive, especially for sections that deviate considerably from standard atlas plates. In these instances, nonlinear adjustments have to be applied manually to match deviations in individual sections. Though this step is time consuming, our results demonstrate that it is important since nonlinear refinement considerably improves the quality of the atlas-registration, as well as the quality of the regional results. Efforts to further automate the atlas-registration step using deep neural networks are underway (DeepSlice)^68^. This QUINT compatible software automates the linear registration step (task currently completed in QuickNII) for whole brain coronal mouse sections, with versions for sagittal and horizontal sections in the pipeline.

The QUINT workflow is a powerful approach for the high-throughput exploration that is needed to unravel the complexity of AD. Using this approach, we further validated the severity of neuroinflammation and pathology accumulation within the brains of aging 5XFAD animals^55, 80–82^ and expanded the extent of anatomical regions investigated in a diverse AD population. AB1-42 levels increased in a widespread manner as 5XFAD mice aged from 6m to 14m. This trend was also seen as near global increases in GFAP and Iba1 were observed across this AD-BXD population^55, 82^. The hippocampus is particularly susceptible to pathology accumulation and atrophy in human patients and a similar decline is also detected in mouse models that display hippocampal degeneration measured via magnetic resonance imaging/IHC^55, 80, 82–90^. We demonstrate that regions that exhibited neurodegeneration, like the Ammon’s horn, were also among those that exhibited the greatest increase in amyloid and neuroinflammation. Previous literature in the 5XFAD model has described visible loss in Layer 5 of the cortex by 9m of age in comparison to Ntg animals^55, 81^, but due to the nature of our current study and the overrepresentation of female 5XFADs we were unable to make this comparison; however, we did detect variation in NeuN load among strains within our AD-BXD population. We can begin to highlight the effect of the 5XFAD transgene and genetic diversity on brain tissue composition in the AD-BXD panel. Genetic differences amongst strains may influence how each strain copes with neuropathology, and the extent of neurodegeneration that occurs with age. Strains can be stratified as resilient or susceptible to AD pathology: with resilient strains potentially mitigating neuron loss in response to neuroinflammation and pathology accumulation, or alternatively staving off severe pathology accumulation all together.

Moreover, we establish an example of how the output from the QUINT workflow can be integrated with a range of data types, including omics data. RNAseq is a common method of profiling gene expression changes between cases and controls and at different disease stages; however, results from bulk tissue samples reflect an average gene expression profile across heterogeneous populations of cells^24^, meaning that expression differences may reflect cell-composition differences across tissue samples, in addition to true transcriptional differences across groups. Determining whether gene hits, established while analyzing bulk RNAseq data, are driven by changes in transcriptional regulation or relative proportions of different cell types in the samples is crucial to establish and properly validate gene candidates of resilience or susceptibility to AD^14, 25, 26^. Recent AD case/control single-nucleus RNA-sequencing datasets offer the opportunity to better resolve such cellular differences^14, 91–94^, but have restrictive technical and cost constraints that can limit the size of such datasets in terms of cells collected and individuals sampled^95^. These limitations as well as the variable performance of deconvolution methods can make it difficult to establish distinct robust cell-type specific differences in gene expression among heterogenous AD populations. While traditional methods for determining cell-type composition, such as IHC or flow cytometry, rely on a limited set of molecular markers and lack in scalability relative to the current rate of data generation, the use of the QUINT workflow can expedite this process. Here we were able to quantify IHC from 39 brains using the QUINT workflow, which streamlined our analysis resulting in high-quality output, and enabled the integration of multiple data types.

To combat the limitations of RNAseq, we integrated IHC-quantified cell composition and RNAseq using mixed modelling correlations. By controlling for age, we were able to establish candidate genes associated with cell composition dependent and independent of the effect of age with AD on variation in load and changes in gene expression. The resulting substantial proportion of genes correlated with load highlights the importance of considering cell composition when analyzing RNAseq data. We also unmasked a unique subset of genes that exhibited no age-related changes in gene expression yet were correlated with variation in load within the age groups examined. Many of the genes that were exclusively significantly correlated with hippocampal formation load following age adjustment were enriched for cell cycle and immune system pathways. By establishing which genes in our dataset are driven by cell and pathology load before and after adjusting for age, we can establish a series of guidelines for prioritizing gene candidates, optimal approaches for modulating genes of interest, and criteria to determine whether candidates should be targeted in a cell-type-specific manner. This study serves as proof-of-concept that IHC data, quantified by the QUINT workflow, can be used as a proxy for cell-type composition in the analysis of RNAseq data, and to demonstrate that changes in gene expression may be relative to variation in cell composition exhibited with age and AD. Due to the nature of this dataset, our analysis was a partial mediation that was only able to begin to disentangle the effect of load, gene expression, and age with AD. Further unravelling this relationship and the effect of the 5XFAD transgene and amyloid accumulation will require additional analyses including nontransgenic animals.

Future investigations will aim to increase the sample size of various AD-BXD strains to confirm and expand upon the current findings. Moreover, the AD-BXD panel has proven to be a strong population to complete genetic mapping of behavioral traits^11, 13–16, 96, 97^, and current efforts are underway to perform genetic mapping of these heritable cell and pathology load traits to identify candidate genes of resilience and susceptibility to AD^98^. These future studies will include non-transgenic littermates, improved intra-strain power by increasing the number of replicates per strain, and the consideration of sex as a biological factor by having equal number of male and female counterparts in each experimental group. Furthermore, this upcoming analysis will utilize the latest version of the CCFv3 (2017) at the intermediate hierarchy established in this study as a baseline for detecting changes in regional cell and pathology load.

In conclusion, we provide the most detailed regional characterization of the 5XFAD mice known to date. The QUINT workflow, with the recent addition of VisuAlign and QCAlign, proved to be a highly effective method and a necessary tool for registering and quantifying cell and pathology changes in diverse disease models like the AD-BXD panel. Achieving high confidence regional output of AD-relevant cell types and pathology also facilitated the exploration of genotype and cell composition relationships. We aim to improve rigor and reproducibility by characterizing the effects of genetic diversity with AD on cell composition and therefore we suggest that bulk-RNAseq data needs to be integrated with cell load to generate robust and reproducible results. By achieving cell and pathology quantification in hemibrains of these mice, we provide a framework for investigators to characterize diverse disease models and integrate their data with a range of behavior and/or omics data.

## Acknowledgements

This study is part of the National Institute on Aging Resilience-AD program and is supported through the NIA parent grant Systems Genetics Analysis of Resilience to Alzheimer’s disease: <u>R01AG057914</u> and supplements R01AG057914-02S1 R01AG057914-03S1 awarded to Dr. Catherine Kaczorowski, The Jackson Laboratory.

The software tools, developed by the Nesys laboratory, University of Oslo, Norway, were funded by EU Horizon 2020, Specific Grant Agreement No. 945539 (Human Brain Project SGA3) awarded to Dr. Jan G. Bjaalie. Workflow optimization for brain-wide spatial analysis to identify regional and cell-type correlates of resilience to Alzheimer’s in the AD-BXD mouse population (BRAINSPACE project) received support from the HBP Voucher Programme call 2019 (ID 66) awarded to Dr. Catherine Kaczorowski, The Jackson Laboratory and Dr. Maja Puchades, University of Oslo.

## Author Contributions

Writing of manuscript: B.G. and S.C.Y.

Revision of manuscript: B.G., S.C.Y., N.H., M.T., K.O., T.M., I.B., H.K., T.B.L., M.A.P, J.G.B., and C.C.K.

Registration and segmentation of brains: B.G.

Curation of registration and segmentation of brains: S.C.Y. and M.A.P.

Generation of results for QUINT workflow validation in QCAlign: B.G., S.C.Y, MT, AO, TO, SS, TM, IB, HK, and US

Data analysis (QCAlign output, Nutil output, IHC-RNAseq integration): B.G., S.C.Y., N.H., and M.T.

Data interpretation: B.G, S.C.Y, N.H, M.T, C.C.K., K.O., M.A.P., J.G.B

Development of experimental design: C.C.K, J.G.B, M.A.P.

Development of QUINT tools (VisuAlign, QCAlign, QNLMask, Nutil): S.C.Y., G.C., N.E.G., T.B.L, M.A.P., J.G.B.

All authors reviewed and approved of the final manuscript.

## Supplemental Information: Figure and Table Legends

**Supplemental Figure 1:** Intermediate hierarchy and QCAlign quality control assessment of atlas registration of thionine sections.

**a.)** Intermediate hierarchy depiction over every thionine section of a representative brain following atlas registration using QuickNII and VisuAlign. Allen Mouse Brain Atlas CCFv3 regions were compiled to make an intermediate hierarchy that promotes the assessment of regional registration. **b.)** Representative quality control assessment of the atlas registration of a thionine slice in QCAlign. Raters assigned grid markers verifying the registration of each point as either accurate, inaccurate, or uncertain.

**Supplemental Figure 2:** QCAlign scores achieved based on quality control assessment of intermediate hierarchy regions.

**a.)** Heatmap of regional accuracy scores per rater per brain. **b)**. Heatmap of regional uncertainty scores per rater per brain. Gray regions were not represented in the brain series and/or did not receive QCAlign scores for the measure. **c.)** Averaged uncertainty scores per intermediate hierarchy region after QuickNII registration alone (white) or after QuickNII and VisuAlign registration (green). Two raters scored the same 5 randomly selected brains after QuickNII registration alone, max n=10 per region (Raters: n= 2 per brain). Up to 10 raters scored the same 5 randomly selected brains after QuickNII and VisuAlign registration, max n=36 per region (Raters: n= 6-10 per brain). Dots represent the mean score across raters per region for 5 brains ±SEM, with the numbers labels representing the number of assessments contributing to each calculation (QuickNII alone labels are below white points, QuickNII + VisuAlign labels are above green points).

**Supplemental Figure 3.** Variation in stain load exists among AD-BXD strains.

Strain averages of **a.)** GFAP, **b.)** Iba1, and **c.)** AB1-42 load across the hippocampal formation and hippocampal intermediate hierarchy subregions. Points are mean load per strain. Each line connects a pair of strain averages across the age groups: 6m and 14m. Only strains with an aged match counterpart are represented (5XFAD mice only, 6m: n=17, 14m: n=18, n= 1-3 per strain). The B6 founder strain is labeled for reference.

**Supplemental Table 1.** Strain, sex, age, 5XFAD genotype, and hemisphere metadata for all animals with IHC completed for this study.

**Supplemental Table 2.** Antibody and dilution information used by NSA for IHC staining.

**Supplemental Table 3.** List of sections removed from individual stain and brain Nutil quantification. Listed sections include those that had greater than 30% damage as measured in QCAlign or were excluded following manual inspection indicating that the majority of the section was distorted and unfit for quantification.

**Supplemental Table 4.** Customized intermediate hierarchy output from QCAlign. List of the 77 intermediate hierarchy regions and the Allen Mouse Brain Atlas IDs that each region is comprised of.

**Supplemental Table 5.** Post-analysis region exclusion parameters. List of 77 regions (compiled by QCAlign from CCFv3 regions) and 5 additional summary regions (Nutil default regions, also compiled from CCFv3 regions) organized by their inclusion or exclusion from QCAlign analysis as represented in figure 2b, Nutil analysis as represented in figure 3a, or IHC and RNAseq integration in figures 4 and 5. “Parent term” are parent IDs, which do not represent any pixels in the CCFv3 and therefore did not generate results; “unassigned pixels” are pixels that are not assigned to a subregion but are instead labeled according to the parent region to which they belong within the Allen Mouse Brain Atlas CCFv3 2015; “low sampling” indicates that less than 20 assessments out of 36 total possible assessments contributed to the mean accuracy QCAlign score for these regions. Some regions were excluded as they had been removed from the brain prior to IHC.

**Supplemental Table 6.** Wilcoxon test results assessing the difference in stain load quantified using QuickNII alone or using QuickNII and VisuAlign for the 55 regions assessed in Figure 2. The regional load per stain per age group among 5XFAD animals was compared between the two methods.

**Supplemental Table 7.** ANOVA results as output from R comparing regional stain load for all intermediate hierarchy regions between 6m and 14m animals. The regional load per stain per age group among 5XFAD animals was compared between the two age groups. FDR-corrected p-values are indicated as FDR_adjusted_pval.

**Supplemental Table 8.** Multilevel correlation results comparing gene expression and hippocampal load correlations both before and after age adjustment for 34 5XFAD animals. FDR-corrected p-values are indicated Age/Non-adjusted p-value (FDR corrected).

## References

1. 2020 Alzheimer’s disease facts and figures. Alzheimer’s & Dementia 16, 391–460 (2020).

2. Congdon, E. E. & Sigurdsson, E. M. Tau-targeting therapies for Alzheimer disease. Nat Rev Neurol 14, 399–415 (2018).

3. Breijyeh, Z. & Karaman, R. Comprehensive Review on Alzheimer’s Disease: Causes and Treatment. Molecules 25, E5789 (2020).

4. Gómez-Tortosa, E. et al. Variability of Age at Onset in Siblings With Familial Alzheimer Disease. Archives of Neurology 64, 1743–1748 (2007).

5. Ryman, D. C. et al. Symptom onset in autosomal dominant Alzheimer disease: A systematic review and meta-analysis. Neurology 83, 253–260 (2014).

6. Dubois, B. et al. Preclinical Alzheimer’s disease: Definition, natural history, and diagnostic criteria. Alzheimers Dement 12, 292–323 (2016).

7. Long, J. M. et al. Preclinical Alzheimer’s disease biomarkers accurately predict cognitive and neuropathological outcomes. Brain 145, 4506–4518 (2022).

8. Chen, Y.-H., Lin, R.-R., Huang, H.-F., Xue, Y.-Y. & Tao, Q.-Q. Microglial Activation, Tau Pathology, and Neurodegeneration Biomarkers Predict Longitudinal Cognitive Decline in Alzheimer’s Disease Continuum. Frontiers in Aging Neuroscience 14, (2022).

9. Moore, S. J., Murphy, G. G. & Cazares, V. A. Turning strains into strengths for understanding psychiatric disorders. Mol Psychiatry 25, 3164–3177 (2020).

10. Onos, K. D., Sukoff Rizzo, S. J., Howell, G. R. & Sasner, M. Toward more predictive genetic mouse models of Alzheimer’s disease. Brain Research Bulletin 122, 1–11 (2016).

11. Neuner, S. M., Heuer, S. E., Huentelman, M. J., O’Connell, K. M. S. & Kaczorowski, C. C. Harnessing Genetic Complexity to Enhance Translatability of Alzheimer’s Disease Mouse Models: A Path toward Precision Medicine. Neuron 101, 399–411.e5 (2019).

12. Neuner, S. M. et al. Translational approaches to understanding resilience to Alzheimer’s disease. Trends in Neurosciences 0, (2022).

13. Heuer, S. E. et al. Identifying the molecular systems that influence cognitive resilience to Alzheimer’s disease in genetically diverse mice. Learn. Mem. 27, 355–371 (2020).

14. Telpoukhovskaia, M. A. et al. Conserved cell-type specific signature of resilience to Alzheimer’s disease nominates role for excitatory cortical neurons. 2022.04.12.487877 Preprint at https://doi.org/10.1101/2022.04.12.487877 (2022).

15. Dai, M. et al. Hypothalamic gene network dysfunction is associated with cognitive decline and body weight loss in Alzheimer’s disease mice. 2022.04.08.487664 Preprint at https://doi.org/10.1101/2022.04.08.487664 (2022).

16. Neuner, S. M., Heuer, S. E., Zhang, J.-G., Philip, V. M. & Kaczorowski, C. C. Identification of Pre-symptomatic Gene Signatures That Predict Resilience to Cognitive Decline in the Genetically Diverse AD-BXD Model. Frontiers in Genetics 10, (2019).

17. Guzmán-Vélez, E. et al. Amyloid-β and tau pathologies relate to distinctive brain dysconnectomics in preclinical autosomal-dominant Alzheimer’s disease. Proceedings of the National Academy of Sciences 119, e2113641119 (2022).

18. Quiroz, Y. T. et al. Association Between Amyloid and Tau Accumulation in Young Adults With Autosomal Dominant Alzheimer Disease. JAMA Neurol 75, 548–556 (2018).

19. Sperling, R. A. et al. Toward defining the preclinical stages of Alzheimer’s disease: recommendations from the National Institute on Aging-Alzheimer’s Association workgroups on diagnostic guidelines for Alzheimer’s disease. Alzheimers Dement 7, 280–292 (2011).

20. Guennewig, B. et al. Defining early changes in Alzheimer’s disease from RNA sequencing of brain regions differentially affected by pathology. Sci Rep 11, 4865 (2021).

21. Hohman, T. J. et al. Asymptomatic Alzheimer disease: Defining resilience. Neurology 87, 2443–2450 (2016).

22. Bocancea, D. I. et al. Measuring Resilience and Resistance in Aging and Alzheimer Disease Using Residual Methods: A Systematic Review and Meta-analysis. Neurology 97, 474–488 (2021).

23. Whitepaper: Defining and investigating cognitive reserve, brain reserve and brain maintenance. Alzheimers Dement 16, 1305–1311 (2020).

24. Li, X. & Wang, C.-Y. From bulk, single-cell to spatial RNA sequencing. Int J Oral Sci 13, 1–6 (2021).

25. Srinivasan, K. et al. Untangling the brain’s neuroinflammatory and neurodegenerative transcriptional responses. Nat Commun 7, 11295 (2016).

26. Consens, M. E. et al. Bulk and Single-Nucleus Transcriptomics Highlight Intra-Telencephalic and Somatostatin Neurons in Alzheimer’s Disease. Frontiers in Molecular Neuroscience 15, (2022).

27. Patrick, E. et al. Deconvolving the contributions of cell-type heterogeneity on cortical gene expression. PLOS Computational Biology 16, e1008120 (2020).

28. Pascal, L. E. et al. Correlation of mRNA and protein levels: Cell type-specific gene expression of cluster designation antigens in the prostate. BMC Genomics 9, 246 (2008).

29. Racle, J. & Gfeller, D. EPIC: A Tool to Estimate the Proportions of Different Cell Types from Bulk Gene Expression Data. Methods Mol Biol 2120, 233–248 (2020).

30. Jew, B. et al. Accurate estimation of cell composition in bulk expression through robust integration of single-cell information. Nat Commun 11, 1971 (2020).

31. Anene, C. A., Taggart, E., Harwood, C. A., Pennington, D. J. & Wang, J. Decosus: An R Framework for Universal Integration of Cell Proportion Estimation Methods. Frontiers in Genetics 13, (2022).

32. Avila Cobos, F., Alquicira-Hernandez, J., Powell, J. E., Mestdagh, P. & De Preter, K. Benchmarking of cell type deconvolution pipelines for transcriptomics data. Nat Commun 11, 5650 (2020).

33. Kang, K., Huang, C., Li, Y., Umbach, D. M. & Li, L. CDSeqR: fast complete deconvolution for gene expression data from bulk tissues. BMC Bioinformatics 22, 262 (2021).

34. Sutton, G. J. et al. Comprehensive evaluation of deconvolution methods for human brain gene expression. Nat Commun 13, 1358 (2022).

35. Doostparast Torshizi, A., Duan, J. & Wang, K. A computational method for direct imputation of cell type-specific expression profiles and cellular compositions from bulk-tissue RNA-Seq in brain disorders. NAR Genomics and Bioinformatics 3, lqab056 (2021).

36. Bjerke, I. E. et al. Data integration through brain atlasing: Human Brain Project tools and strategies. Eur Psychiatry 50, 70–76 (2018).

37. Boline, J., Lee, E.-F. & Toga, A. Digital atlases as a framework for data sharing. Frontiers in Neuroscience 2, (2008).

38. Yates, S. C. et al. QUINT: Workflow for Quantification and Spatial Analysis of Features in Histological Images From Rodent Brain. Frontiers in Neuroinformatics 13, (2019).

39. Puchades, M. A., Csucs, G., Ledergerber, D., Leergaard, T. B. & Bjaalie, J. G. Spatial registration of serial microscopic brain images to three-dimensional reference atlases with the QuickNII tool. PLOS ONE 14, e0216796 (2019).

40. Berg, S. et al. ilastik: interactive machine learning for (bio)image analysis. Nat Methods 16, 1226–1232 (2019).

41. Groeneboom, N. E., Yates, S. C., Puchades, M. A. & Bjaalie, J. G. Nutil: A Pre- and Post- processing Toolbox for Histological Rodent Brain Section Images. Frontiers in Neuroinformatics 14, (2020).

42. Hammelrath, L. et al. Morphological maturation of the mouse brain: An in vivo MRI and histology investigation. NeuroImage 125, 144–152 (2016).

43. Hikishima, K. et al. In vivo microscopic voxel-based morphometry with a brain template to characterize strain-specific structures in the mouse brain. Sci Rep 7, 85 (2017).

44. Wang, N. et al. Variability and heritability of mouse brain structure: Microscopic MRI atlases and connectomes for diverse strains. Neuroimage 222, 117274 (2020).

45. Purger, D. et al. A histology-based atlas of the C57BL/6J mouse brain deformably registered to in vivo MRI for localized radiation and surgical targeting. Phys Med Biol 54, 7315–7327 (2009).

46. Chen, X. J. et al. Neuroanatomical differences between mouse strains as shown by high-resolution 3D MRI. Neuroimage 29, 99–105 (2006).

47. Wang, Q. et al. The Allen Mouse Brain Common Coordinate Framework: A 3D Reference Atlas. Cell 181, 936–953.e20 (2020).

48. Schindelin, J., et al. Fiji: an open-source platform for biological-image analysis. Nat Methods 9, 676–682 (2012).

49. Love, M. I., Huber, W. & Anders, S. Moderated estimation of fold change and dispersion for RNA-seq data with DESeq2. Genome Biology 15, 550 (2014).

50. Makowski, D., Ben-Shachar, M., Patil, I. & Lüdecke, D. Methods and Algorithms for Correlation Analysis in R. JOSS 5, 2306 (2020).

51. Liao, Y., Wang, J., Jaehnig, E. J., Shi, Z. & Zhang, B. WebGestalt 2019: gene set analysis toolkit with revamped UIs and APIs. Nucleic Acids Research 47, W199–W205 (2019).

52. Wang, J., Vasaikar, S., Shi, Z., Greer, M. & Zhang, B. WebGestalt 2017: a more comprehensive, powerful, flexible and interactive gene set enrichment analysis toolkit. Nucleic Acids Research 45, W130–W137 (2017).

53. Wang, J., Duncan, D., Shi, Z. & Zhang, B. WEB-based GEne SeT AnaLysis Toolkit (WebGestalt): update 2013. Nucleic Acids Research 41, W77–W83 (2013).

54. Zhang, B., Kirov, S. & Snoddy, J. WebGestalt: an integrated system for exploring gene sets in various biological contexts. Nucleic Acids Res 33, W741–748 (2005).

55. Oakley, H., et al. Intraneuronal beta-Amyloid Aggregates, Neurodegeneration, and Neuron Loss in Transgenic Mice with Five Familial Alzheimer’s Disease Mutations: Potential Factors in Amyloid Plaque Formation. Journal of Neuroscience 26, 10129–10140 (2006).

56. Tsui, K. C. et al. Distribution and inter-regional relationship of amyloid-beta plaque deposition in a 5xFAD mouse model of Alzheimer’s disease. Frontiers in Aging Neuroscience 14, (2022).

57. Rao, Y. L. et al. Hippocampus and its involvement in Alzheimer’s disease: a review. 3 Biotech 12, 55 (2022).

58. Habes, M. et al. Disentangling Heterogeneity in Alzheimer’s Disease and Related Dementias Using Data-Driven Methods. Biol Psychiatry 88, 70–82 (2020).

59. Altschuler, S. J. & Wu, L. F. Cellular heterogeneity: when do differences make a difference? Cell 141, 559–563 (2010).

60. Wang, X. et al. Deciphering cellular transcriptional alterations in Alzheimer’s disease brains. Molecular Neurodegeneration 15, 38 (2020).

61. Akasaka-Manya, K. & Manya, H. The Role of APP O-Glycosylation in Alzheimer’s Disease. Biomolecules 10, 1569 (2020).

62. Akasaka-Manya, K. et al. Excess APP O-glycosylation by GalNAc-T6 decreases Aβ production. The Journal of Biochemistry 161, 99–111 (2017).

63. Zhang, L. et al. Sex-specific DNA methylation differences in Alzheimer’s disease pathology. Acta Neuropathol Commun 9, 77 (2021).

64. Jordan-Sciutto, K. L., Malaiyandi, L. M. & Bowser, R. Altered Distribution of Cell Cycle Transcriptional Regulators during Alzheimer Disease. J Neuropathol Exp Neurol 61, 358– 367 (2002).

65. Barbash, S. et al. Alzheimer’s brains show inter-related changes in RNA and lipid metabolism. Neurobiology of Disease 106, 1–13 (2017).

66. Liu, E. Y., Cali, C. P. & Lee, E. B. RNA metabolism in neurodegenerative disease. Dis Model Mech 10, 509–518 (2017).

67. Lauridsen, K. et al. A Semi-Automated Workflow for Brain Slice Histology Alignment, Registration, and Cell Quantification (SHARCQ). eNeuro 9, (2022).

68. Carey, H. et al. DeepSlice: rapid fully automatic registration of mouse brain imaging to a volumetric atlas. 2022.04.28.489953 Preprint at https://doi.org/10.1101/2022.04.28.489953 (2022).

69. Ju, T. et al. 3D volume reconstruction of a mouse brain from histological sections using warp filtering. Journal of Neuroscience Methods 156, 84–100 (2006).

70. Wahlsten, D., Hudspeth, W. J. & Bernhardt, K. Implications of genetic variation in mouse brain structure for electrode placement by stereotaxic surgery. J. Comp. Neurol. 162, 519– 531 (1975).

71. Johnson, G. A. et al. HiDiver: A Suite of Methods to Merge Magnetic Resonance Histology, Light Sheet Microscopy, and Complete Brain Delineations. 2022.02.10.479607 Preprint at https://doi.org/10.1101/2022.02.10.479607 (2022).

72. Johnson, G. A. et al. Waxholm Space: An image-based reference for coordinating mouse brain research. NeuroImage 53, 365–372 (2010).

73. Takata, N., Sato, N., Komaki, Y., Okano, H. & Tanaka, K. F. Flexible annotation atlas of the mouse brain: combining and dividing brain structures of the Allen Brain Atlas while maintaining anatomical hierarchy. Sci Rep 11, 6234 (2021).

74. Ni, H. et al. DeepMapi: a Fully Automatic Registration Method for Mesoscopic Optical Brain Images Using Convolutional Neural Networks. Neuroinform 19, 267–284 (2021).

75. Fürth, D. et al. An interactive framework for whole-brain maps at cellular resolution. Nat Neurosci 21, 139–149 (2018).

76. Lein, E. S. et al. Genome-wide atlas of gene expression in the adult mouse brain. Nature 445, 168–176 (2007).

77. Osten, P. & Margrie, T. W. Mapping brain circuitry with a light microscope. Nature Methods 10, 515–523 (2013).

78. Gurdon, B. et al. Brain-wide spatial analysis to identify region-specific changes in cell composition associated with resilience to Alzheimer’s disease in the AD-BXD mouse population. Alzheimer’s & Dementia 16, e047613 (2020).

79. Gurdon, B. & Kaczorowski, C. Pursuit of precision medicine: Systems biology approaches in Alzheimer’s disease mouse models. Neurobiol Dis 161, 105558 (2021).

80. Jawhar, S., Trawicka, A., Jenneckens, C., Bayer, T. A. & Wirths, O. Motor deficits, neuron loss, and reduced anxiety coinciding with axonal degeneration and intraneuronal Aβ aggregation in the 5XFAD mouse model of Alzheimer’s disease. Neurobiology of Aging 33, 196.e29–196.e40 (2012).

81. Eimer, W. A. & Vassar, R. Neuron loss in the 5XFAD mouse model of Alzheimer’s disease correlates with intraneuronal Aβ42 accumulation and Caspase-3 activation. Molecular Neurodegeneration 8, 2 (2013).

82. Forner, S. et al. Systematic phenotyping and characterization of the 5xFAD mouse model of Alzheimer’s disease. Sci Data 8, 270 (2021).

83. Oblak, A. L. et al. Comprehensive Evaluation of the 5XFAD Mouse Model for Preclinical Testing Applications: A MODEL-AD Study. Front Aging Neurosci 13, 713726 (2021).

84. Frankó, E., Joly, O. & Initiative, for the A. D. N. Evaluating Alzheimer’s Disease Progression Using Rate of Regional Hippocampal Atrophy. PLOS ONE 8, e71354 (2013).

85. Apostolova, L. G. et al. Subregional hippocampal atrophy predicts Alzheimer’s dementia in the cognitively normal. Neurobiology of Aging 31, 1077–1088 (2010).

86. Uysal, G. & Ozturk, M. Hippocampal atrophy based Alzheimer’s disease diagnosis via machine learning methods. Journal of Neuroscience Methods 337, 108669 (2020).

87. Mueller, S. G. et al. Hippocampal atrophy patterns in mild cognitive impairment and Alzheimer’s disease. Human Brain Mapping 31, 1339–1347 (2010).

88. DeTure, M. A. & Dickson, D. W. The neuropathological diagnosis of Alzheimer’s disease. Molecular Neurodegeneration 14, 32 (2019).

89. Coupé, P., Manjón, J. V., Lanuza, E. & Catheline, G. Lifespan Changes of the Human Brain In Alzheimer’s Disease. Sci Rep 9, 3998 (2019).

90. Braak, H., Alafuzoff, I., Arzberger, T., Kretzschmar, H. & Del Tredici, K. Staging of Alzheimer disease-associated neurofibrillary pathology using paraffin sections and immunocytochemistry. Acta Neuropathol 112, 389–404 (2006).

91. Mathys, H. et al. Single-cell transcriptomic analysis of Alzheimer’s disease. Nature 570, 332–337 (2019).

92. Cain, A. et al. Multi-cellular communities are perturbed in the aging human brain and Alzheimer’s disease. 2020.12.22.424084 Preprint at https://doi.org/10.1101/2020.12.22.424084 (2022).

93. Zhou, Y. et al. Human and mouse single-nucleus transcriptomics reveal TREM2-dependent and TREM2-independent cellular responses in Alzheimer’s disease. Nat Med 26, 131–142 (2020).

94. Leng, K. et al. Molecular characterization of selectively vulnerable neurons in Alzheimer’s disease. Nat Neurosci 24, 276–287 (2021).

95. Park, Y. et al. Single-cell deconvolution of 3,000 post-mortem brain samples for eQTL and GWAS dissection in mental disorders. http://biorxiv.org/lookup/doi/10.1101/2021.01.21.426000 (2021) doi:10.1101/2021.01.21.426000.

96. Neuner, S. M. et al. TRPC3 channels critically regulate hippocampal excitability and contextual fear memory. Behav Brain Res 281, 69–77 (2015).

97. Neuner, S. M. et al. Systems genetics identifies Hp1bp3 as a novel modulator of cognitive aging. Neurobiology of Aging 46, 58–67 (2016).

98. Gurdon, B. et al. Brain-wide spatial analysis reveals cell-type-specific genetic modifiers of Alzheimer’s disease progression. Alzheimer’s & Dementia 18, e061853 (2022).

