## Supplementary figures and images for "Detecting the effect of genetic diversity on brain composition in an Alzheimer’s disease mouse model"

### Supplemental Figures

Supplemental Table 1

a

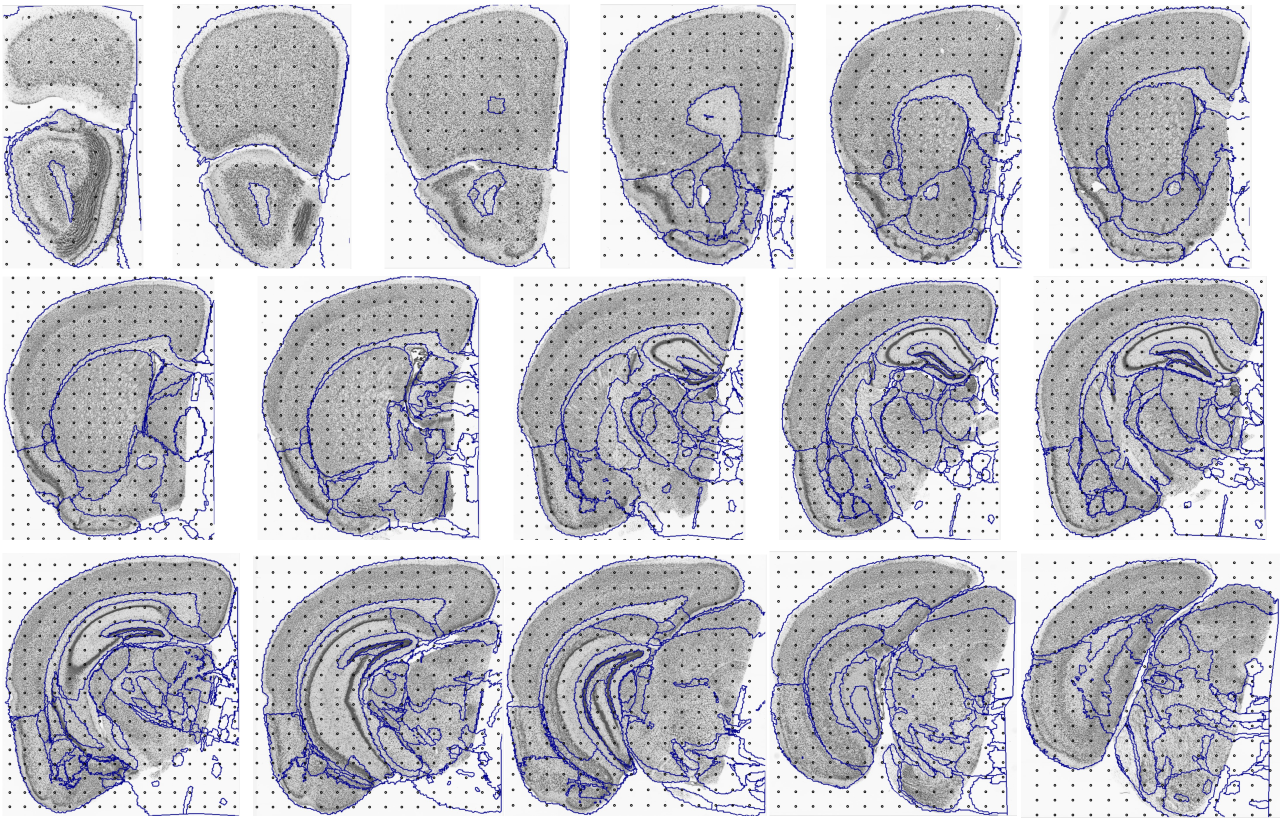

b

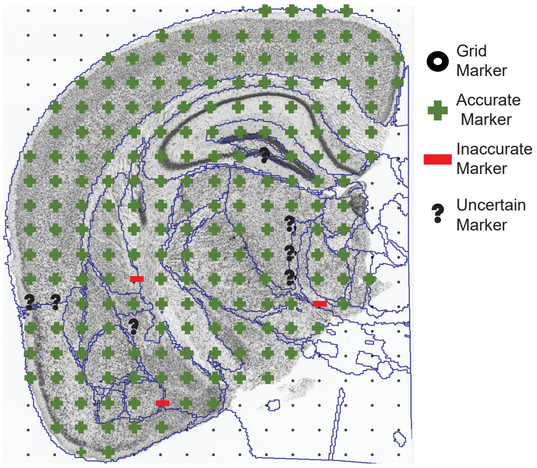

Supplemental Figure 2

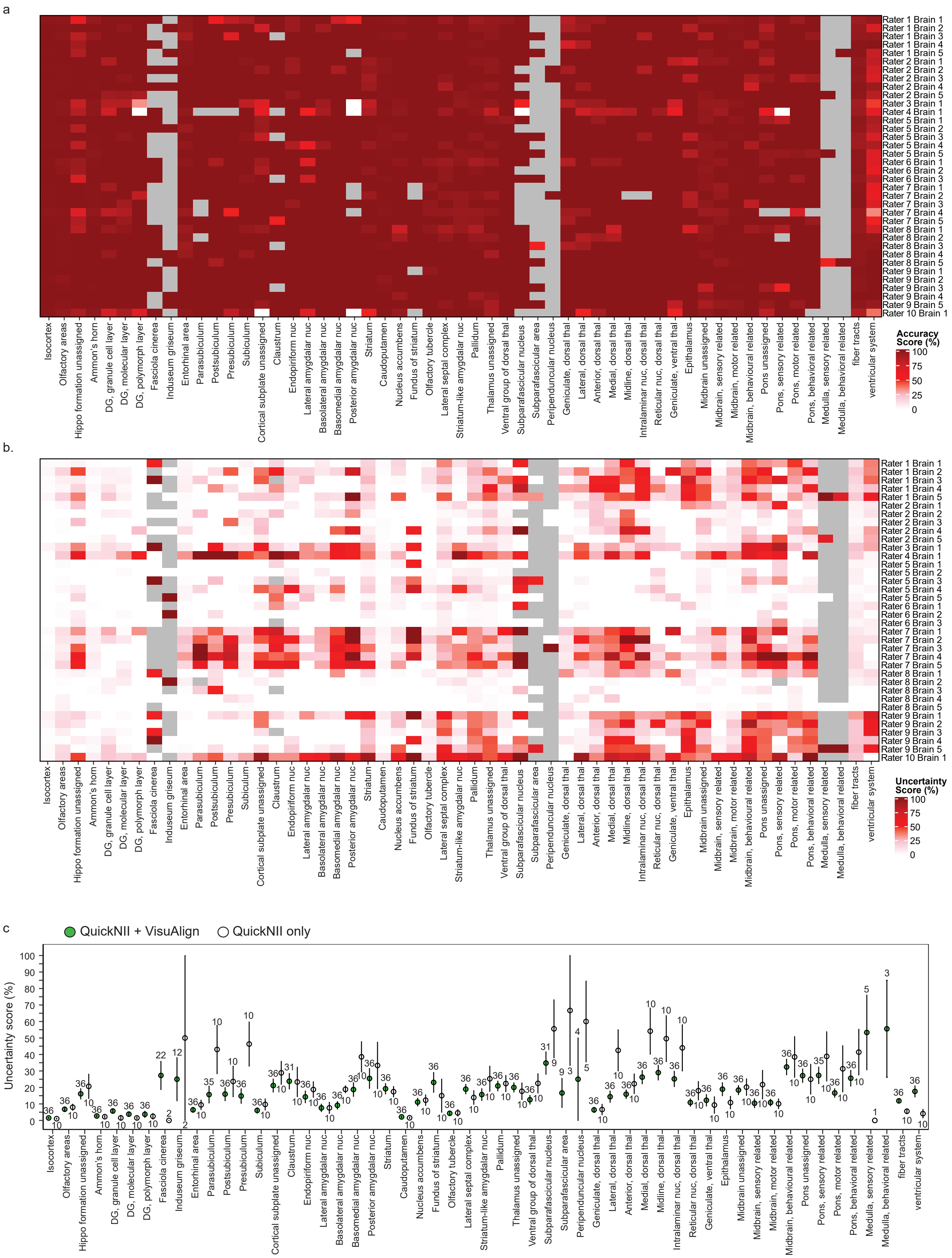

Supplemental Figure 3

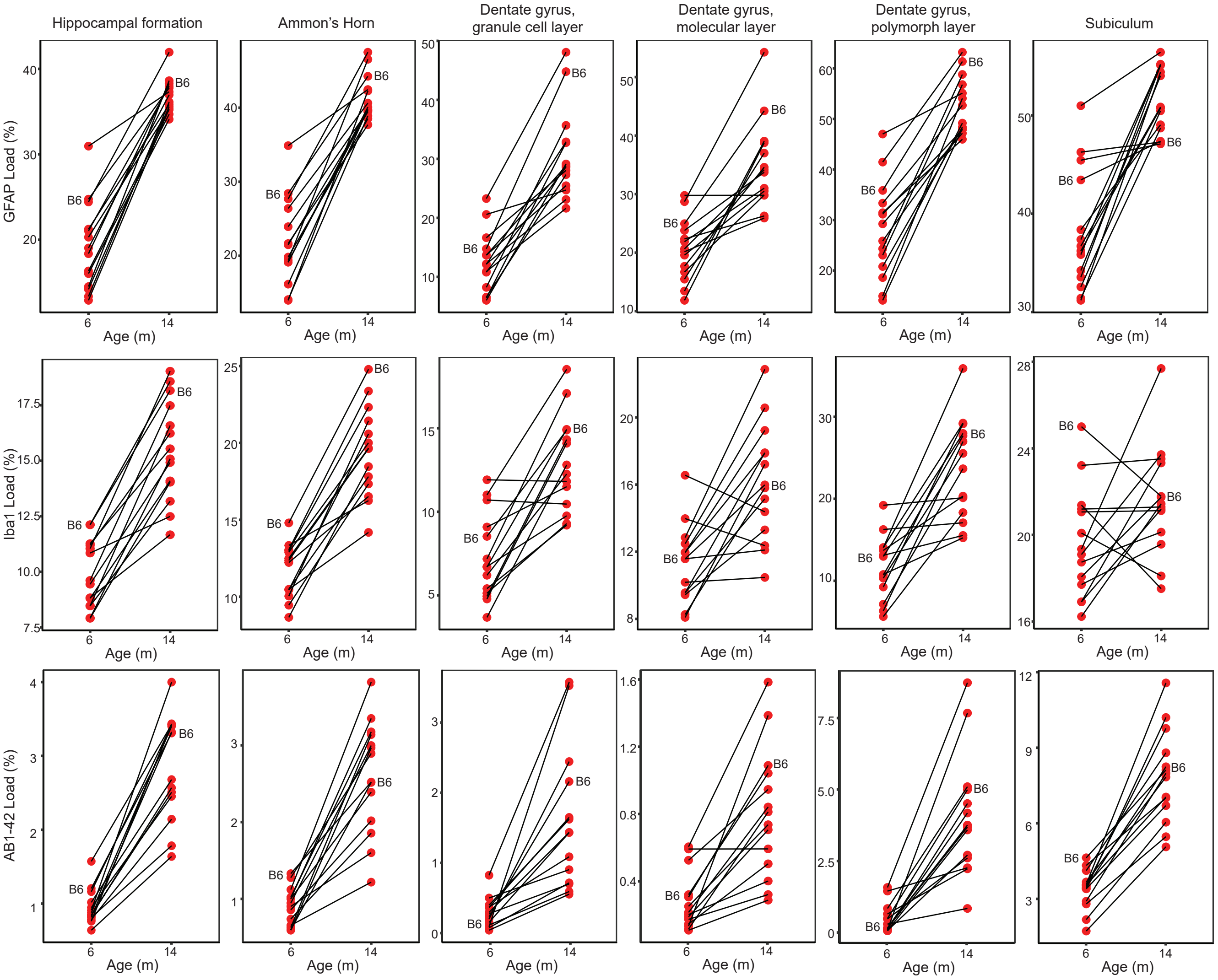
